# Extracting multi-way chromatin contacts from Hi-C data

**DOI:** 10.1101/2021.11.04.467227

**Authors:** Lei Liu, Bokai Zhang, Changbong Hyeon

## Abstract

There is a growing realization that multi-way chromatin contacts formed in chromosome structures are fundamental units of gene regulation. However, due to the paucity and complexity of such contacts, it is challenging to detect and identify them using experiments. Based on an assumption that chromosome structures can be mapped onto a network of Gaussian polymer, here we derive analytic expressions for *n*-body contact probabilities (*n* > 2) among chromatin loci based on pairwise genomic contact frequencies available in Hi-C, and show that multi-way contact probability maps can in principle be extracted from Hi-C. The three-body (triplet) contact probabilities, calculated from our theory, are in good correlation with those from measurements including Tri-C, MC-4C and SPRITE. Maps of multi-way chromatin contacts calculated from our analytic expressions can not only complement experimental measurements, but also can offer better understanding of the related issues, such as cell-line dependent assemblies of multiple genes and enhancers to chromatin hubs, competition between long-range and short-range multi-way contacts, and condensates of multiple CTCF anchors.

**Author summary:** The importance of DNA looping is often mentioned as the initiation step of gene expression. However, there are growing evidences that ‘chromatin hubs’ comprised of multiple genes and enhancers play vital roles in gene expressions and regulations. Currently a number of experimental techniques to detect and identify multi-way chromosome interactions are available; yet detection of such multi-body interactions is statistically challenging. This study proposes a method to predict multi-way chromatin contacts from pair-wise contact frequencies available in Hi-C dataset. Since chromosomes are made of polymer chains, the pairwise contact probabilities are not entirely independent from each other, but certain types of correlations are present reflecting the underlying chromosome structure. We extract these correlations hidden in Hi-C dataset by leveraging theoretical argument based on polymer physics.

## Introduction

Recent advances in experimental techniques [1–6] offer unprecedented glimpses into the chromosome structures inside cell nuclei, in the form of pairwise distances and contact frequencies between genomic loci. Gene expressions are, however, realized when multiple genomic loci, e.g., promoters and enhancers separated over large genomic distances, are brought together to form a regulatory element [7–10], which underscores the importance of resolving chromatin interactions beyond pairwise two-body contacts. In particular, the transcriptional regulation is fine-tuned via a complex network of cooperative and competitive interactions among nearby genes mediated by transcription factors [11, 12].

A plethora of experimental methods have recently been developed to detect multi-way contacts in chromosome [13], which include super-resolution chromatin tracing [14], and both ligation-based (3way-4C [15], COLA [16], Tri-C [17], MC-4C [18, 19], Pore-C [20] and single-cell techniques [21–24]) and ligation-free methods (GAM [25] and SPRITE [26]). Detection of multi-way chromatin contacts using these experimental methods, however, are statistically limited due to the paucity of such contacts. The probability of detecting a particular *n*-body contacts from a genomic region of interest consisting of *N* statistical segments is approximately 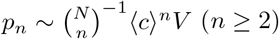, where ⟨*c*⟩ ~ *N/V* is the effective concentration of the segments in the volume *V* which can be approximated using the Flory radius [27] (*R*_*F*_ ~ *N*^*ν*^) to 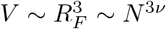 with *ν*(≥ 1/3) being the Flory exponent [28, 29]. Since 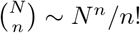 for *N* ≫ *n*, it is expected that the *n*-body contact probability scales as *p*_*n*_ ~ *n*!*/N* ^3*ν*(*n*−1)^, which makes experimental detection of *n*-body contacts with larger *n* statistically more demanding. To get around the detection problem, one usually either conducts a genome-wide study at low resolution (a small *N*), or performs a high-resolution experiment by focusing only on the contacts formed at a few prescribed sites [16, 30].

Since the polymer physics idea was first used to explore the physical characteristics of chromosomes as long polymer chains confined in a small nuclear space [28, 31–38], computational strategies of incorporating genomic constraints from experimental measurements and epigenomic information into polymer-based modeling of 3D chromosome structure have recently gained much traction [39–54]. Many studies, which generate an ensemble of 3D chromosome structures, highlight the heterogeneous and probabilistic nature of chromosome structure [45, 47, 50, 51, 54]. Once an ensemble of structural models of chromosomes are obtained from computational approaches, it is straightforward to count the multi-way chromatin contacts directly from them and to quantify the corresponding contact probabilities. Specifically, recent studies, which have generated 3D chromosome structures based on the strings and binders switch (SBS) model, have shown that the probabilities of triplet chromatin contacts for HoxD and *α*-globin regions calculated from the structures are in good agreement with those from 3way-4C and Tri-C experiments, respectively [55, 56]. CHROMatin mIXture (CHROMATIX) model was used to address the functional relevance of many-body contacts among genomic elements enriched at transcriptionally active loci [30].

In this study, we calculate *n*-body contact probabilities from Hi-C data by using analytic expressions derived from the formalism of Heterogeneous Loop Model (HLM) [50, 51, 57]. Although the original aim of HLM was to reconstruct 3D chromosome structures from Hi-C [50, 51], it is still possible to calculate the map of *n*-body contact probabilities without explicitly counting those contacts from the generated chromosome structures. Through comparisons between the multi-way chromatin contacts derived from HLM and those from three separate measurements from Tri-C, MC-4C, and SPRITE, we will show that the *n*-body contact probability maps are in good agreement with experimental measurements (**Results**). We also explore the relation between pairwise and higher-order contacts and discuss how to avoid the most evident false-positiveness in the analysis of multi-way chromatin contacts experiments (**Discussions**). Finally, the numerical details of training HLM based on Hi-C and the mathematical derivations for contact probabilities are provided in the **Methods** section.

## Results

### Polymer theory of *n*-body chromatin contacts

The chromatin fiber in a genomic region of interest is modeled as a coarse-grained polymer chain composed of *N* monomers (or sites), each representing a chromatin segment of a prescribed genomic length. We assume that the chromatin effective energy landscape can be described by a summation of harmonic restraints on the spatial distances between all monomer pairs,

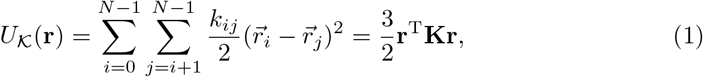

where 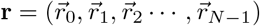 specifies the 3D structure of the polymer chain, and 𝒦 is a *N*-by-*N* symmetric stiffness matrix of elements *k*_*ij*_. The so-called Kirchhoff matrix is defined by **K** ≡ 𝒟 − 𝒦, where 𝒟 is a diagonal matrix with elements 𝒟_*ii*_ = ∑_*j*_ *k*_*ij*_. We assume that the probability of the chromatin to a adopt a particular structure is given by

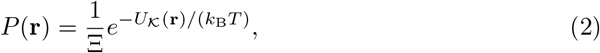

where *k*_B_*T* is our energy unit (the Boltzmann contant times the temperature) and Ξ = (det(**K**)/(2*π*)^*N*^)^3/2^ is a normalization constant such that the integration of Eq. 2 over all possible structures equals to 1.

The primary assumption of HLM (Eq. 1) results in a probability distribution of the physical distance between any (*i, j*) monomers, *r*_*ij*_ [50, 58]

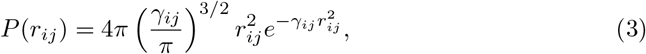

where *γ*_*ij*_ is a function of **K**-matrix with a positive value. More specifically, *γ*_*ij*_ = (*σ*_*ii*_ + *σ*_*jj*_ − 2*σ*_*ij*_)^−1^ /2 for *i* > 0 and 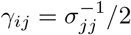 for *i* = 0, in which *σ*_*ij*_ denotes the (*i, j*)-th element of the inverse matrix of **K**. To assess our assumption, we compare Eq. 3 with the fluorescence *in situ* hybridization (FISH) data recently reported by Takei *et al* [24]. By using DNA seqFISH+ method, they measured the 3D coordinates of 2,460 loci spaced approximately 1 Mb apart across the whole genome, together with additional 60 consecutive loci at 25 kb resolution on each chromosome, in 446 mouse embryonic stem cells (Fig 1A).

**Fig 1.**
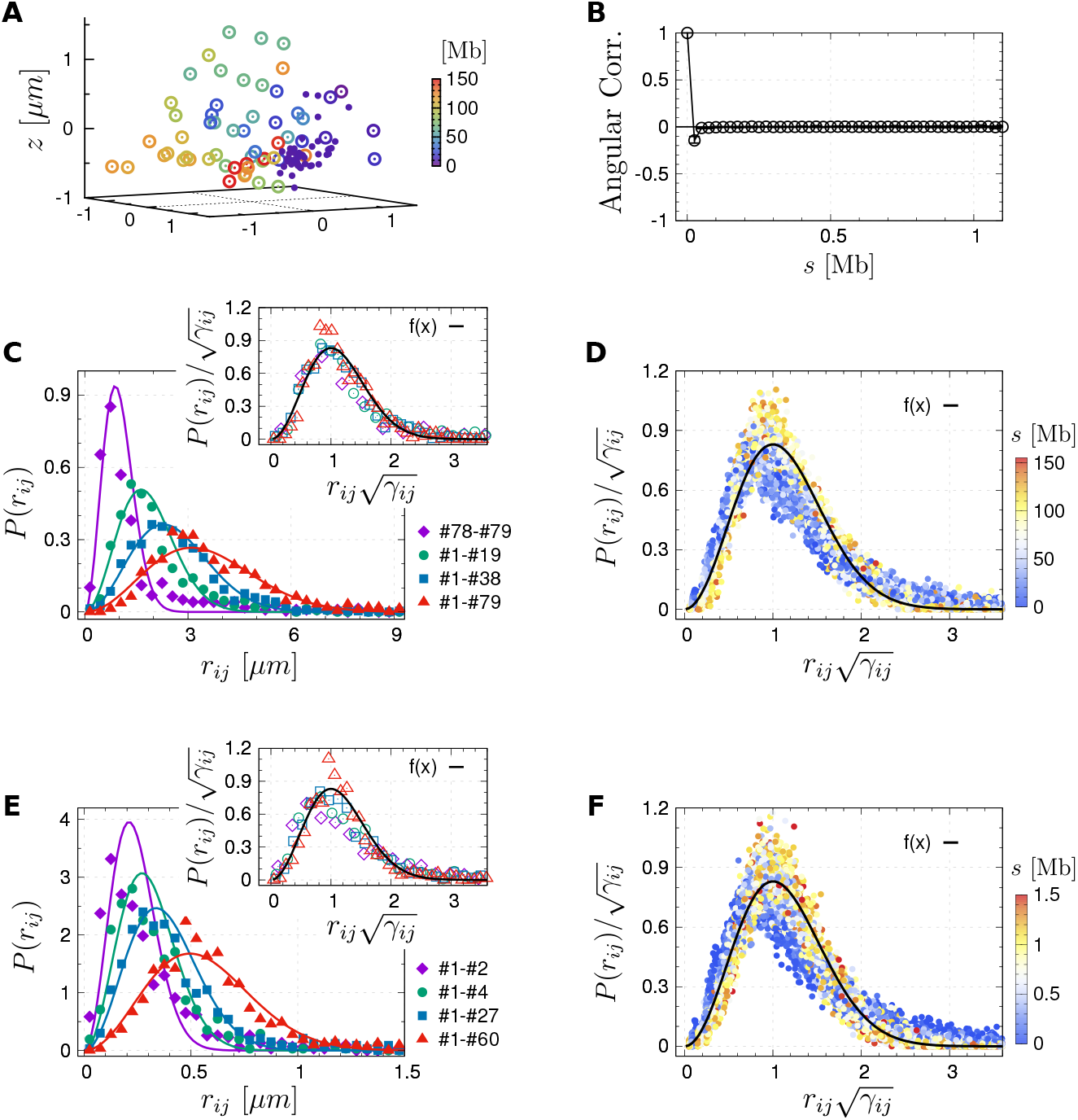
Chain organization of chr3 of mouse ES cells measured in the DNA seqFISH+ experiment [24]. **(A)** 3D positions of 151 imaged loci spaced about 1 Mb apart (large empty dots) and 60 consecutive loci with an equal space of 25 kb (small solid dots) in one of 446 sample cells. **(B)** Angular correlation of the chromatin segments at 25 kb resolution. **(C)** Distribution of the physical distance of four loci pairs at 1 Mb resolution (markers) with their fittings to our theory (lines, Eq. 3), which are rescaled by using the fitting parameter (*γ*_*ij*_) and plotted in the inset. **(D)** Rescaled pairwise distance distributions of all loci pairs, which are colored by their genomic lengths. **(E, F)** Similar as **(C, D)** but at 25 kb resolution.

At 1 Mb resolution (Fig 1C), the distribution of the physical distances between four loci pairs on chromosome 3 (distinguished by markers of different colors) can be well fitted by Eq. 3. When we replot the experimental data with a scaled distance 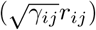, all rescaled data are distributed around a master curve

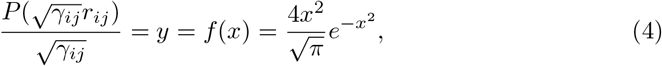

which is shown as the black solid line in the inset of Fig 1C. Taking all 151 imaged loci on chr3 as a test case, we analyzed the distance distributions of all intra-chromosome loci pairs (i.e., 151 × 150/2 = 11, 325 pairs), calculated *γ* for each pair, and plotted the rescaled data. As shown in Fig 1D, regardless of the genomic distance between these loci (labeled by different colors of the dots), data collected over the whole chromosome lie around the master curve, lending support to the validity of Eq 3. This also holds when analyzing the distance distribution of loci at a finer resolution of 25 kb (Fig 1E and F).

Next, the cross-linking probability function *F*(*r*), describes how likely two chromatin fragments are captured by cross-linking agents if they are spaced by a distance of *r*. Based on *F*(*r*), one can count the contact frequency for any monomer pair in a given structural ensemble, as well as the higher-order contact frequency among any group of *n* monomers (2 *< n* ≤ *N*). The higher-order *n*-body contact is formed when *n* monomers (genomic segments) are simultaneously in spatial proximity and within a capture radius of cross-linking agent. The *n*-body contact probability, e.g., 3-body (*n* = 3, triplet) contact probability between *i, j*, and *k* monomer, *p*_*ijk*_, can thus be calculated by integrating the probability of a particular chromatin structure (Eq. 2) multiplied by the chance to form simultaneous cross-linking among the *n* monomers in that structure over all possible chromatin structures. Thanks to the special form of the energy function (Eq. 1), HLM yields analytic formulae. Specifically, when the effective capture radius of the cross-linking agent is denoted by *r*_*c*_, we obtain the following results:

- The pairwise contact probability *p*_*ij*_ can be written as

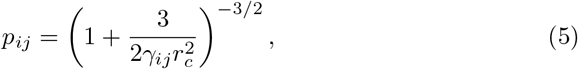

with *γ*_*ij*_ defined in Eq. 3.
- The triplet contact probability *p*_*ijk*_ is

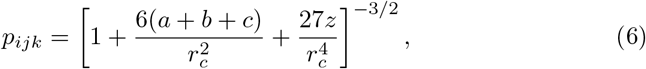

where *a* = *σ*_*ii*_ + *σ*_*jj*_ − 2*σ*_*ij*_, *b* = *σ*_*ij*_ + *σ*_*jk*_ − *σ*_*ik*_ − *σ*_*jj*_, *c* = *σ*_*jj*_ + *σ*_*kk*_ − 2*σ*_*jk*_, and *z* = *ac* − *b*^2^.
- The 4-body contact probability *p*_*ijkl*_ is

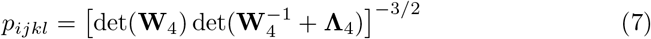

where **W**_4_ is a 3 × 3 matrix of the form 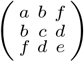 with *d* = *σ*_*jk*_ + *σ*_*kl*_ − *σ*_*jl*_ − *σ*_*kk*_, *e* = *σ*_*kk*_ + *σ*_*ll*_ − 2*σ*_*kl*_, *f* = *σ*_*ik*_ + *σ*_*jl*_ − *σ*_*il*_ − *σ*_*jk*_, and 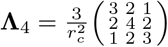. The parameters *a, b, c* are identical to those in Eq.6.

Readers are referred to the **Methods** section for detailed derivations. As briely descried above, these results are exactly equivalent to reconstructing an ensemble of 3D chromatin structures followed by counting explicitly *n*-body contact frequencies in the ensemble.

The stiffness matrix 𝒦 has to be determined before calculating the *n*-body contact probability and predict specific multi-way chromatin interactions. As illustrated in Fig 2, 𝒦-matrix can be determined from 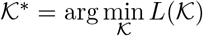, i.e., by minimizing *L*(𝒦) – a cost function that quantifies the difference between the 𝒦-dependent pairwise contact probabilities of the model (*p*_*ij*_(𝒦)) and those from pairwise contacts experiment 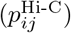 (see the **Methods** section for the details). We next compare our predictions of many-body chromatin contacts with those from measurements found in the literature. Detailed information on which genomic region are modeled, the cell types, the model resolutions and the experiment sources are provided in S1 Table.

**Fig 2.**
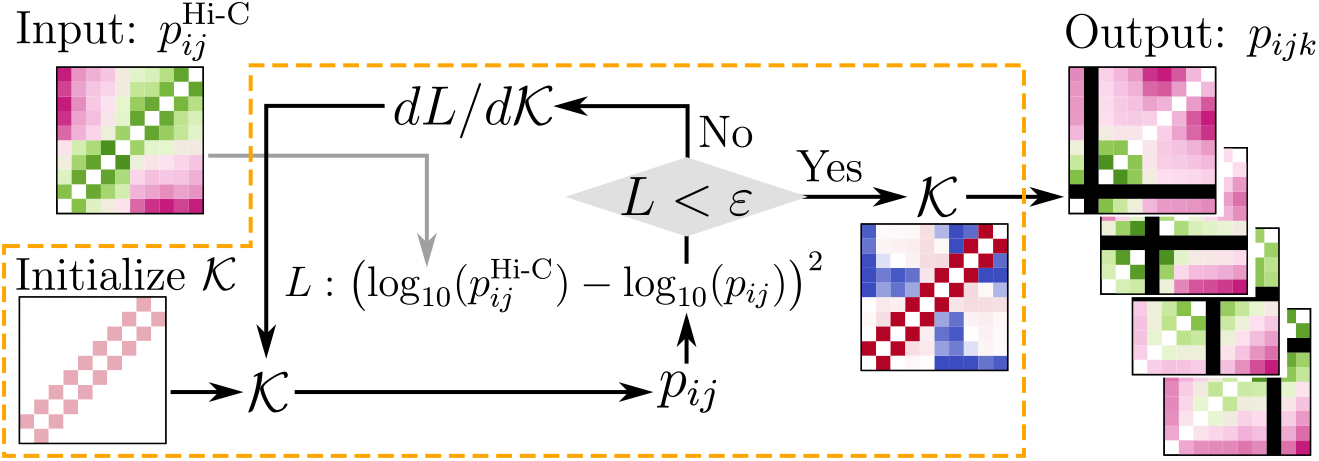
Flow chart of HLM that is enclosed by the dashed orange box. It takes two-body contacts from Hi-C as input, and calculates *n*-body (*n* > 2) contact probability. The stiffness matrix 𝒦 is updated until the algorithm finds 𝒦* that minimizes the cost function *L* = *L*(𝒦) which quantifies the difference between pairwise contact probabilities of Hi-C data and those determined from 𝒦 matrix.

### Comparison with Tri-C

Our first case study for our HLM-based theoretical predictions (see Eq.6) concerns the mouse *α*-globin locus, an extensively studied model system [61], whose three-body contacts are available from Tri-C measurement [17]. The globin genes are flanked by several CTCF-binding sites (the 3rd row in Fig 3A) in both erythroid and embryonic stem (ES) cells. The Capture-C heatmaps of the two cell types (Fig 3B. The top and bottom panels are for the ES and erythroid cells, respectively) show that the erythroid cell display more frequent contacts (stronger interactions) between the *α*-globin genes (Hba-a1/2) and the five upstream enhancer elements (R1, R2, R3, R4 and Rm).

**Fig 3.**
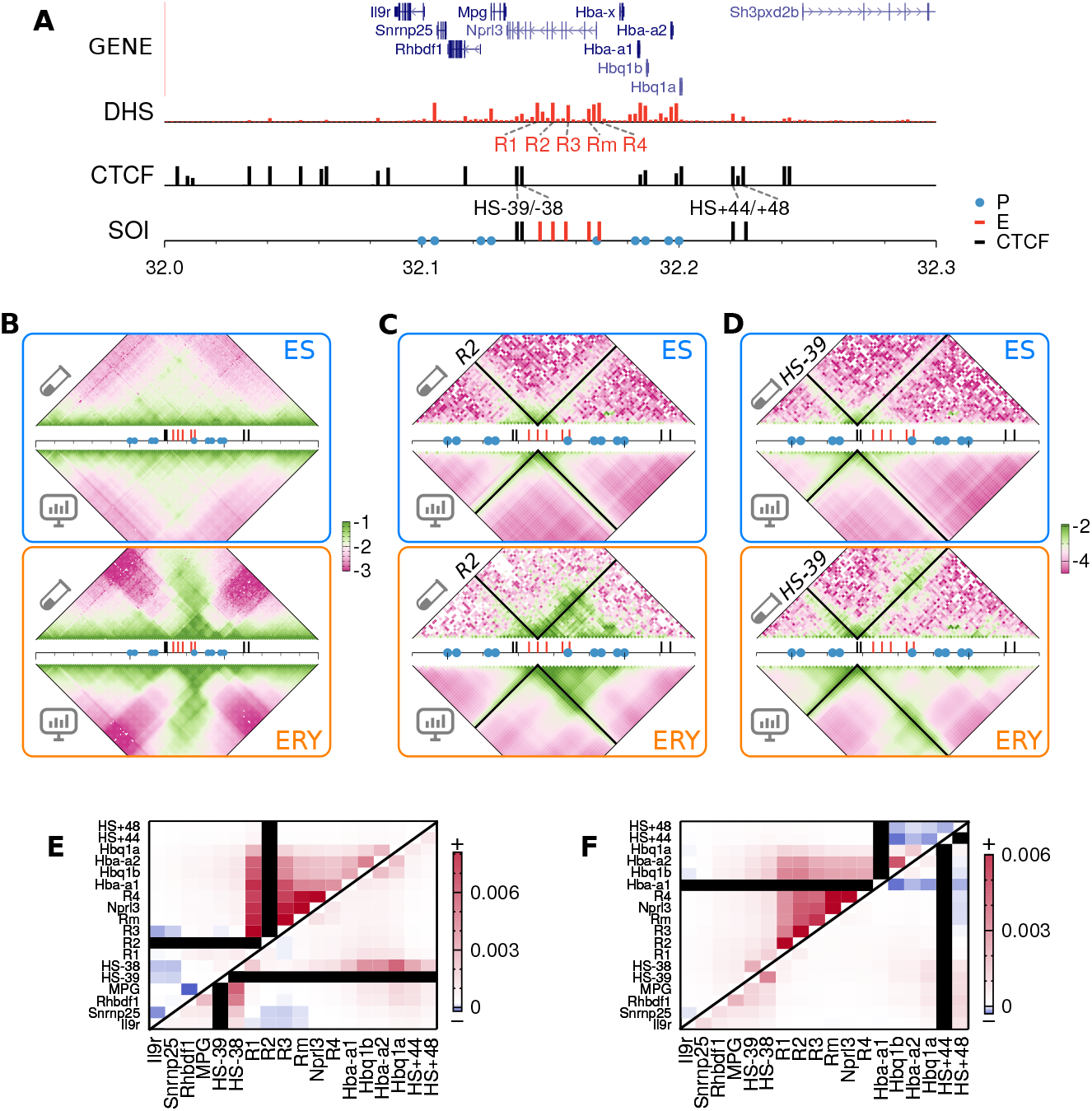
Three-body contacts predicted with HLM versus those from Tri-C. **(A)** Gene notation, DNase hypersensitive sites, CTCF-binding sites [59, 60] and genomic regions of interest (sites of interest, SOI) at mouse *α*-globin locus on chr11. **(B)** log_10_(*p*_*ij*_) from Capture-C (top triangle) and HLM (bottom triangle) at 2-kb resolution. **(C, D)** log_10_(*p*_*ijk*_) from Tri-C (top) and HLM (bottom) at *k* = R2 and *k* = HS-39, whose positions are labeled with black strips. The Pearson correlations between Tri-C and HLM are 0.80 (ES) and 0.75 (erythroid) for *k* = R2, and 0.89 (EC) and 0.85 (erythroid) for *k* = HS-39. **(E)** Cell-line difference of the predicted three-body contacts, 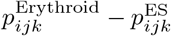, among the SOIs with respect to the viewpoints R2, HS-39 and **(F)** Hba-a1, HS+44.

The 𝒦-matrices of HLM, obtained separately for both cell lines based on Hi-C data, are not only used to calculate *p*_*ij*_ that captures the variation of local pairwise contacts upon ES-to-erythroid cell differentiation (Fig 3B), but also to predict triplet contacts (*p*_*ijk*_) at different genomic locus in a cell-type specific manner (Fig 3C and D). We calculated the two maps of triplet contact probability (i) at the genomic locus of the strongest enhancer R2 (Fig 3C) and (ii) at the upstream CTCF-binding site HS-39 (Fig 3D). The Pearson correlations (PC) between the HLM-predicted triplet contacts and those from Tri-C data are generally high (PC ≈ 0.75 − 0.89. See the caption of Fig 3 for details). In particular, for the triplet contact probability map calculated at the HS-39 viewpoint, the correlations are comparable to those reported by a simulation study using the SBS model (PC ≈ 0.8) [56]. We further calculated three correlation coefficients specifically designed to compare contact matrices, namely the distance-corrected Pearson correlation [46], stratum-adjusted correlation [62] and stratified Pearson correlation (S1 Fig). As summarized in S2 Table, overall, HLM matches better with Tri-C experiment than SBS model.

The comparison between triplet contact probabilities in ES cells over all sites of interest (SOIs) and those in erythroid cells underscores the change in higher-order contacts upon cell differentiation (Fig 3E). Upon the activation of *α*-globin gene (ES → erythroid cell), R2 simultaneously interacts with the gene promoters and its nearby enhancers especially R1 more frequently, which suggests the formation of a regulatory hub. HS-39, HS-38 and distal downstream CTCF binding sites are not involved in this hub, but form diffuse interactions in between. These findings from the HLM-predicted triplet contacts are consistent with the Tri-C analyses conducted at R2 and HS-39 [17] (see S2 Fig).

We made additional comparisons at the globin gene promoter Hba-a1 and the downstream CTCF boundary element HS+44, whose triplet contact probabilities have not yet been measured (Fig 3F). At the viewpoint Hba-a1, the promoter interacts with upstream enhancers in a cooperative manner, which supports the above analyses. By contrast, in the presence of contact with a downstream boundary element, Hba-a1 becomes less likely to interact with another downstream boundary site in erythroid cells.

### Comparison with multi-contact 4C sequencing (MC-4C)

Next, we study the formation of higher-order contacts among CTCF binding sites in the cohesin release factor (WAPL) lacking cells [63]. According to the loop extrusion model [64–66], the ring-shaped protein complex, cohesin, binds to DNA and progressively enlarges chromatin loop until the dynamics is hindered by two convergently orientated CTCF-bound sites. Knockout of the WAPL promotes the chromatin loop extension [63].

We carried out HLM calculation for a 1.28-Mb genomic region on chromosome 8, which contains 13 CTCF-binding sites, for both wild-type (WT) and WAPL-deficient (ΔWAPL) human chronic myeloid leukemia (HAP1) cells, whose triplet contacts were measured with MC-4C [18]. Although the Hi-C dataset (Fig 4A, top) is noisier than that used in the first case study (Fig 3B, top), the HLM-predicted triplet contacts at the CTCF binding sites E (Fig 4B) and K (Fig 4C) show patterns similar to those from the MC-4C measurement [18]. To account for the enrichment of long-range triplet contacts comprised of distal CTCF binding elements (e.g., the triplet A, H and K) in the WAPL-lacking cells, Allahyar *et al*. [18] put forward a “traffic jam” of CTCF roadblock-trapped cohesin complexes.

**Fig 4.**
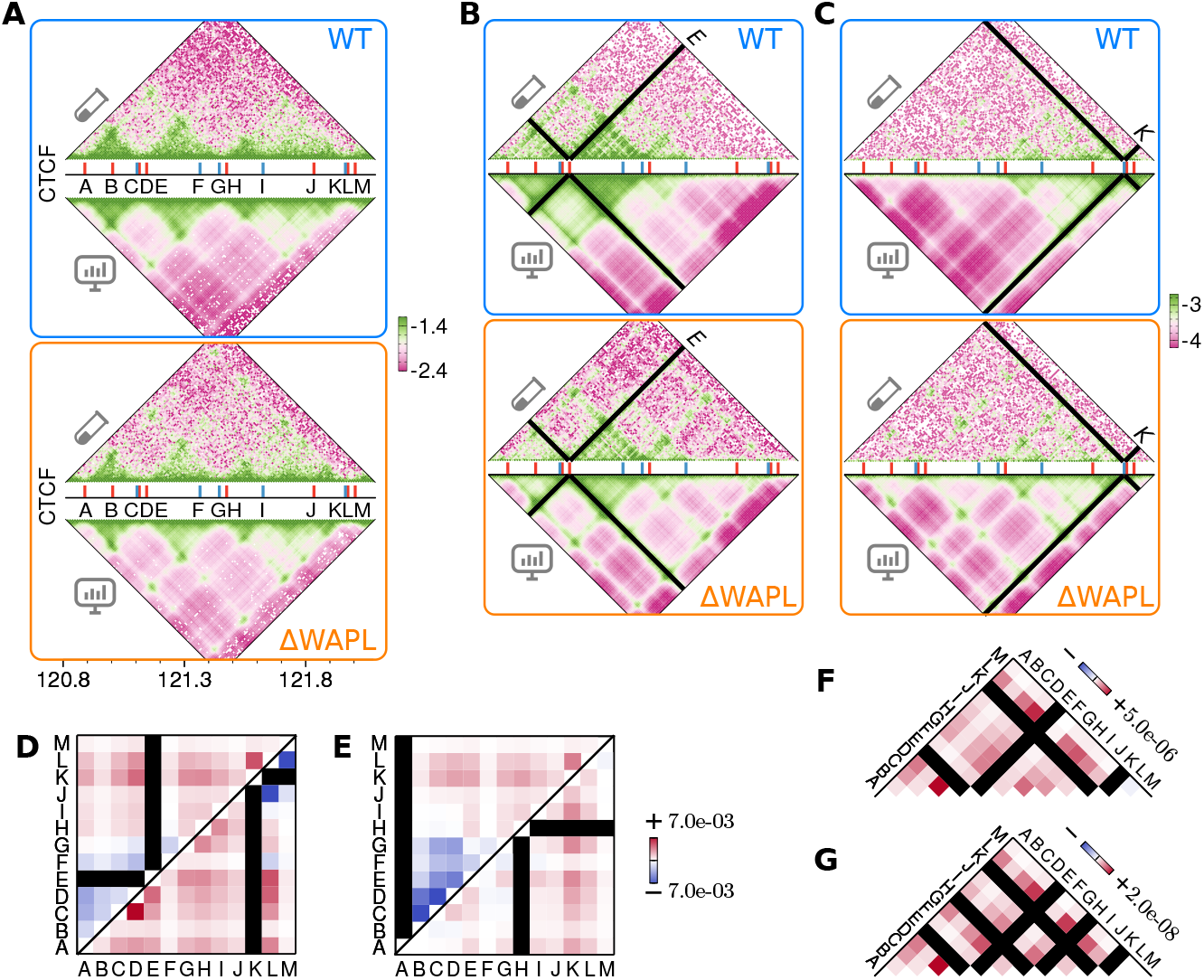
Comparing many-body contact predictions with MC-4C. **(A)** Comparing log_10_(*p*_*ij*_) from Hi-C and HLM at 10-kb resolution of a 1.28 Mb region on chr8, where there are thirteen CTCF-binding sites labeled from A to M. The genomic position of forward- and backward-oriented CTCF sites are marked with red and blue sticks, respectively. **(B, C)** Comparing log_10_(*p*_*ijk*_) from MC-4C and HLM at a viewpoint of site E and K, respectively. From the viewpoint of E (K), the values of PC between the model and the MC-4C experiment are 0.83 (0.81) and 0.71 (0.73) for the WT and ΔWAPL cells, respectively. **(D)** Cell-line difference of the predicted three-body contacts, namely 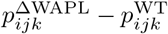, among the CTCF-binding sites from the viewpoint of A, E and **(E)** H, K, respectively. **(F, G)** Cell-line difference of four- and five-body contacts from double- and triple-anchored viewpoints, whose positions are marked with the black strips. The numbers label the maximum amplitudes of changes.

Examining the WAPL knockout-induced change in triplet contacts among CTCF binding sites, we plot the changes of 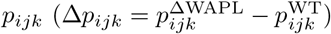 at four viewpoints, *k* = E, K and A, H in Fig 4D and E, respectively. While the site A is colocalized more frequently with the distant downstream sites K and L in ΔWAPL cell, its interaction with the nearby loci (sites B-G) is reduced significantly (e.g., *p*_ABC_ is reduced by more than 60% from 1.1 × 10^−3^ to 4.2 × 10^−4^). Similar patterns of “long-range enrichment and short-range depletion of triplet contacts” are identified for those from other upstream and downstream viewpoints (e.g., the sites E and K). Although the MC-4C measurement [18] did not highlight the short-range depletion of triplet contacts, our finding is consistent with the contact frequencies measured with respect to the sites E and K (S3 Fig A). In fact, the two Hi-C datasets, one for WT and the other for ΔWAPL cell, show that the pairwise contact frequencies of ΔWAPL cells in the domains flanked by short-range CTCF binding sites are smaller than those of the WT (S3 Fig B), which directly confirm the short-range depletion of contacts. The observation of long-range enrichment and short-range depletion of triplet contacts does not always hold; the central sites H and I are the two exceptions that do not demonstrate depletions of short-range triplet contacts.

The aggregation of cohesin molecules has been identified in ΔWAPL cells with super-resolution imaging [18]; however, it has not been known whether or not a similar type of aggregation occurs in CTCF-bound chromatin sites. To explore the possibility of condensation of CTCF-bound chromatin sites, we calculate higher-order chromatin contacts in both conditions of WT and ΔWAPL cells. The changes in 4-body and 5-body contacts among CTCF-bound sites from WT to ΔWAPL cells from double- and triple-anchored view points are all positive, namely, 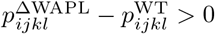 (Fig 4F) and 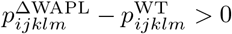 (Fig 4G), which lends supports to the picture surmising the clustering of domain boundary in ΔWAPL cells. Although their probabilities are small, up to 4-fold increases of 4-body and 5-body contacts are identified among the SOIs (see S4 Fig).

### Comparison with SPRITE

Lastly, we model a 0.45 Mb genomic region of GM12878 cell (Fig 5A and B), which contains transcriptionally active genes BCL2 and multiple super-enhancers, annotated in Ref. [30] by Perez-Rathke *et al*., and compared the predicted triplet contacts with the results from SPRITE [26]. The human BCL2 gene encodes a membrane protein which blocks the apoptotic death in B-lymphoblastoid cells.

**Fig 5.**
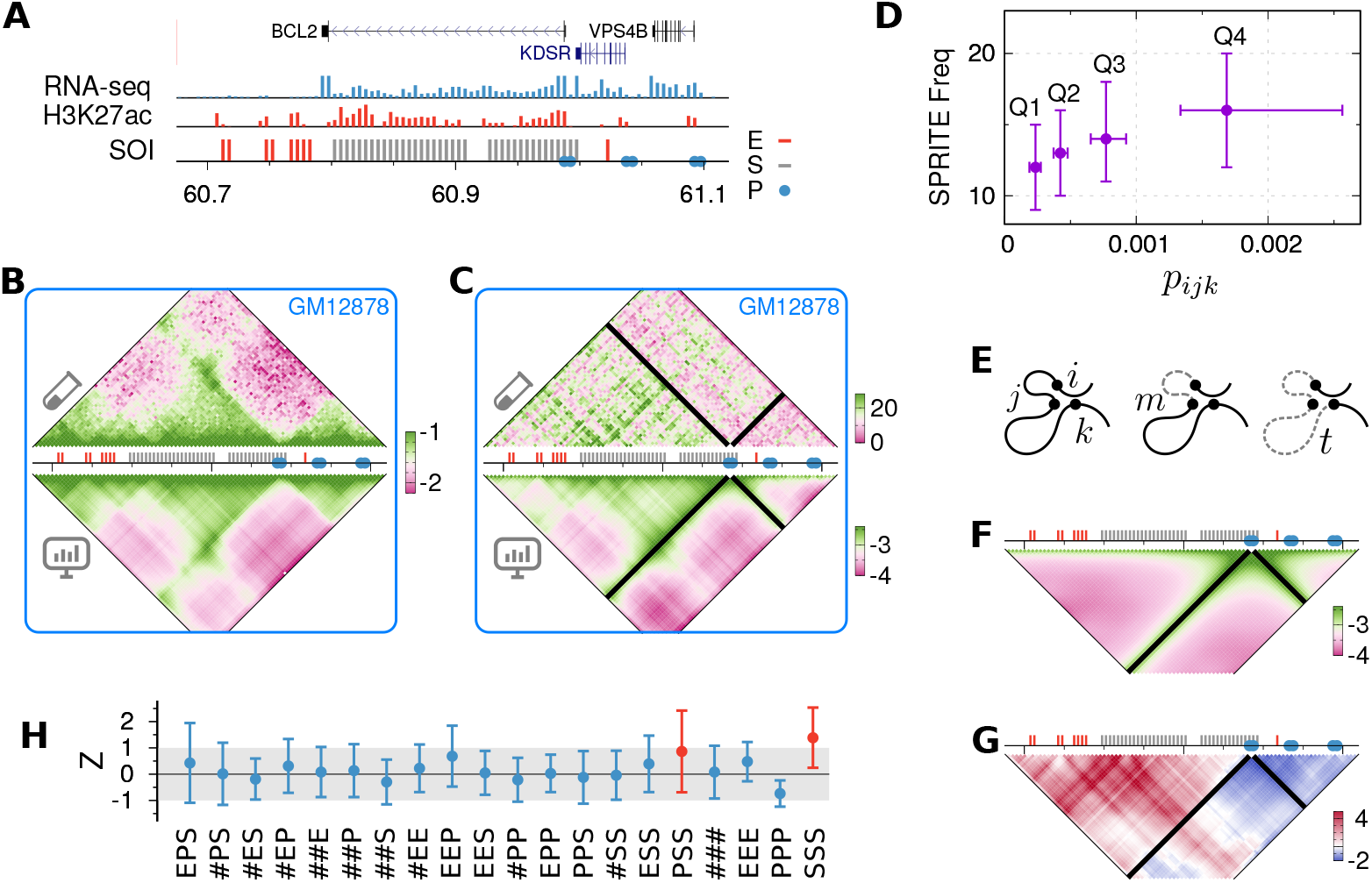
Comparing triplet contact predictions with SPRITE. **(A)** Genes, RNA-seq and H3K27ac ChIP-seq signals [60, 67] in a 0.45 Mb region on human chr18 with the positions of three types of SOIs (Enhancers, Promoters and Super-enhancers). **(B)** log_10_(*p*_*ij*_) from Hi-C compared with that from HLM at 5-kb resolution. **(C)** The triplet contact frequency from SPRITE is compared with log_10_(*p*_*ijk*_) from HLM anchored at an active promoter site. **(D)** SPRITE frequency of triplets which are divided into four quantiles based on ascending order of *p*_*ijk*_. **(E)** For three sites, *i < j < k*, along a polymer chain, the minor and major sections denoted by *m* and *t*, respectively, are depicted with dashed lines. **(F)** The expected three-body contact probability 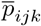, and **(G)** the specificity Z-score 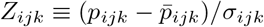. **(H)** The mean specificity Z-score of triplet contacts with respect to all possible combinations of annotations (E: Enhancers, P: Promoters, S: Super-enhancers, #: sites without any annotation). The error bars are the standard deviations.

We expect that the similarity between the model and the experiment is not so high as the foregoing two case studies, because the data from SPRITE, designed for a genome-wide study, is sparse with 5-kb resolution, and the chromatin fragments cocaptured by SPRITE are not necessarily in spatial proximity since SPRITE does not distinguish between direct and indirect cross-linkings. Indeed, a weak correlation is found in the triplet contacts anchored at the active BCL2 gene promoter (PC = 0.18; see Fig 5C). To validate our models against the measurements, we sort the triplets in an ascending order of their predicted contact probabilities, and divide them into four quantiles. Then the frequency of the quantilized triplets in SPRITE increases with the predicted probability, *p*_*ijk*_ (Fig 5D).

To identify the specificity in a single cell type, we defined an expected triplet contact probability 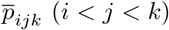 as the mean contact probability of all triplets in the “sections” of interest along the chain (Fig 5E). The minor and major sections are determined as *m* = min(|*j* − *i*|, |*k* − *j*|) and *t* = |*k* − *i* |, respectively. 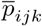 depends only on *m* and *t* (Fig 5F), which reflects the chain connectivity and the global compactness of chromatin. The ratio of the difference between the observed and expected probabilities scaled by the standard deviation is then defined as the *specificity Z-score*, 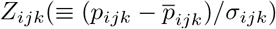, for each triplet (Fig 5G). The triplet contacts are deemed specific if *Z*_*ijk*_ > 1.

*Z*_*ijk*_ averaged over all possible combinations of functional annotations in the region of interest are shown in Fig 5H. The concurrent triplet contacts between two super-enhancers with a promoter (PSS) or with a third super-enhancer (SSS) are specifically enriched at this loci. This finding comports well with the Perez-Rathke *et al* ‘s finding that multiple super-enhancers are involved in higher-order chromatin contacts more frequently than other elements [30]. We confirm similar results from modeling at another active locus (S5 Fig).

## Discussions

### Relation between pairwise and triplet contacts

For a given polymer chain, is there any simple relation to associate two-body contacts with three-body contacts? Through both experimental and theoretical studies, there has been much effort to address this question and related issues in the context of concurrent chromatin interactions [14, 15, 18, 19, 68].

Here, we address this issue in a principled way by considering a Gaussian polymer chain without any specific interaction as a reference system. In the heatmaps of *n*-body contact probability of the Gaussian polymer chain consisting of *N* monomers, the enrichments of many-body contacts are observed near the diagonal part of *N* × *N* matrix as well as around the viewpoints (Fig 6A-C). For any triplet, *ijk*(*i < j < k*), the contact probability reads (see Eq. 26 for the derivation)

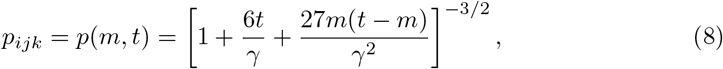

where *m* = min(|*j* − *i* |, |*k* − *j*|) is the minor section, *t* = |*k* − *i*| is the major section of the triplet (Fig 5E), and *γ* is the stiffness constant which restrains the consecutive sites. For *t* ≫ *m*, it follows that *p*_*ijk*_ ~ *t*^−3/2^, the scaling exponent of which is identical to that of the two-body contact probability [28, 29]. For a fixed *t, p*(*m, t*) changes non-monotonically as a function of *m*, having its minimum and maximum at *m* = *t/*2 and *m* = 1, respectively (Fig 6B). According to Eq. 8 the specificity Z-score defined in the subsection **Comparison with SPRITE** satisfies *Z*_*ijk*_ = 0 for all triplets in a Gaussian chain, as anticipated.

**Fig 6.**
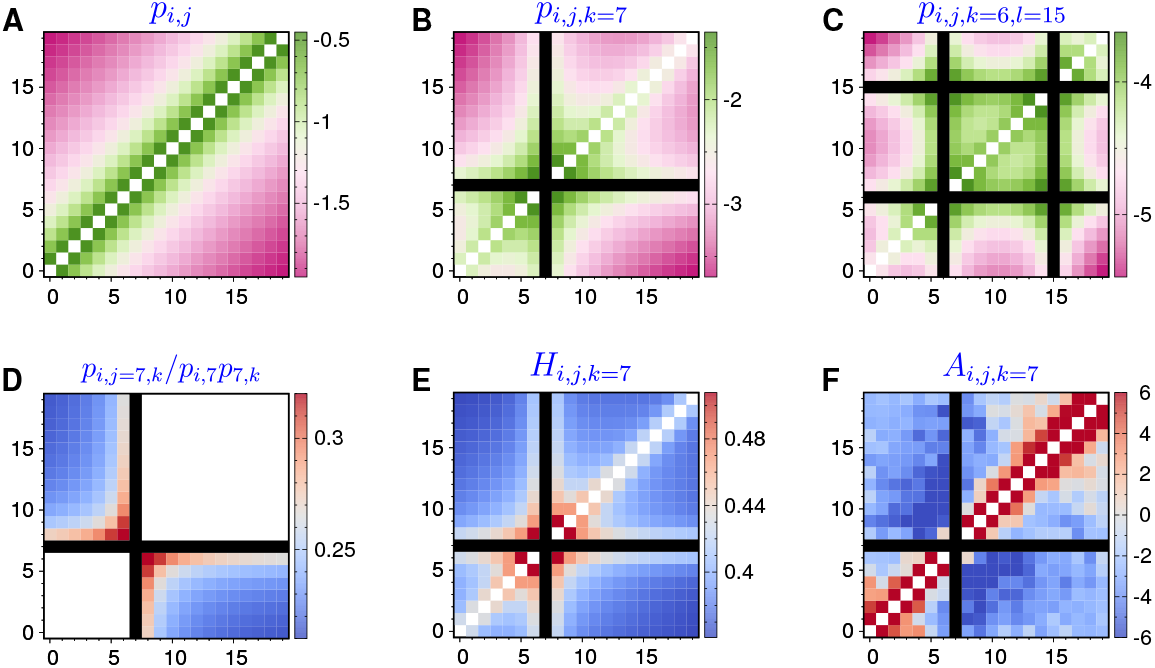
Many-body contacts in a Gaussian polymer chain consisting of 20 sites. **(A)** Two-body contact probability log_10_(*p*_*ij*_). **(B)** Three-body contact probability log_10_(*p*_*ijk*_) anchored at the 7-th site (*k* = 7). **(C)** Four-body contact probability log_10_(*p*_*ijkl*_) double anchored at the 6-th and 15-th sites (*k* = 6, *l* = 15). **(D-F)** Boost factor *p*_*ijk*_*/p*_*ij*_*p*_*jk*_, hub-score *H*_*ijk*_, and association z-score *A*_*ijk*_ from the viewpoint of the 7-th site.

An enhancement of triplet contact compared to two independent pairwise loops was discussed by Polovnikov *et al*. [68], who used the delta function for the cross-linking probability between genomic sites, such that sites were considered in contact only if they overlapped with each other. It was found that *p*_*ijk*_*/p*_*ij*_*p*_*jk*_ ≥ 1 with a maximum at *m* = *t/*2. This finding, however, is not universally held if one considers other types of cross-linking probability. When Gaussian and Heaviside step functions are used as the cross-linking probabilities, triplet contact probabilities are always suppressed in comparison with the product of two binary contact probabilities, i.e., *p*_*ijk*_*/p*_*ij*_*p*_*jk*_ ≤ 1, and the largest suppression occurs at *m* = *t/*2 (Fig 6D and S6 Fig D).

No clear-cut relation exists between pairwise (*p*_*ij*_) and triplet contact probabilities (*p*_*ijk*_), which however was the underlying presumption of the data processing in several experiments. For example, a hub score was calculated to identify synergistic interaction hubs in the analysis of 3way-4C data [15], which was defined by *H*_*ijk*_ = 3*p*_*ijk*_/(*p*_*ij*_*p*_*jk*_ + *p*_*ij*_*p*_*ik*_ + *p*_*jk*_*p*_*ik*_) approximately, whereas our Gaussian chain model clarifies that the hub score is not a constant (Fig 6E). In fact, as discussed in the experiment [15], enrichment of the score is always found at the triplets close to any viewpoint (e.g., Fig 3 and 4 in Ref. [15]).

To detect cooperative (competitive) contacts, *an association Z-score*, defined as 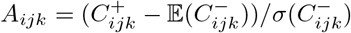, were computed in the analyses of MC-4C measurements [18, 19]. 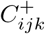 is the number of reads cocapturing the genetic sites *i, j* and *k* (the viewpoint). 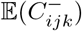 and 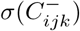 are the mean and standard deviation of 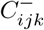, the number of reads capturing the sites *j* and *k* but not *i*. A positive (negative) *A*_*ijk*_, which implies that the chance of site *j* being in association with *k* and any other third site is (dis)favored when site *i* is interacting with *k*, was interpreted as a measure of cooperative (competitive) triplet contacts. However, the calculation based on Gaussian polymer chain shows that adjacent monomer pairs along the chain display a greater association z-scores *A*_*ijk*_ (Fig 6F). This issue of false-positiveness in the short-range cooperativity among the multiple sites has been noticed in Ref. [18].

A recent super-resolution imaging study on chromatin fibers in IMR90 cells has explored whether or not the association between two genetic loci facilitates the contact with a third site [14]. For triplets *ijk* satisfying *i < j < k*, two conditional probabilities 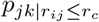 and 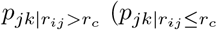 is the contact probability between the *k*-th and *j*-th sites under the condition that *i*-th and *j*-th sites are in contact, and 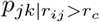 is similarly defined) were compared with the unconditioned contact probability *p*_*jk*_. It was found that regardless of cell types and other chemical treatments, a relation 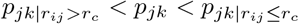 was always satisfied among three probabilities, for most triplets of genomic loci [14]. Our theoretical framework allows us to validate this relation for any triplet in a polymer chain (see S7 Fig). The generic characteristics of multi-body contacts of Gaussian chain mentioned above do not depend on the specific form of cross-linking probability (S6 Fig).

As a further test, we counted the average, maximum, and minimum pairwise contact probabilities of the triplets anchored at R2 in the *α*-globin locus of mouse erythroid cells (S8 Fig B-D). They have dissimilar patterns with respect to the results from Tri-C experiment (S8 Fig A), which is also reflected by their overall lower Pearson correlations than HLM calculated at two different viewpoints in two different cell types (S8 Fig E). The relation between pairwise and multi-way contacts is more complicated than intuition.

### Miscellaneous

The prediction of HLM is not always consistent with experiments. For example, when scrutinizing the WAPL knockout-induced change of triplet contacts at viewpoint of the CTCF binding site E, an enrichment involving site G can be found in MC-4C (see the left panel of S3 Fig) which is not in agreement with our model prediction (Fig 4D). Many factors can contribute to this discordance, such as the quality of the Hi-C dataset at the locus of interest. Despite much effort devoted in processing raw Hi-C data, bias which remains in the Hi-C contact matrix after normalization will propagate to the model and subsequent predictions on higher-order contacts.

Another apparent missing component of our approach is the excluded volume interaction, which plays an indisputable role in determining polymer behavior. However, as shown in Fig 1, chromatin segment pairs have non-zero probabilities at small physical distances, in contrast to the expectation that segments are not allowed to overlap with each other. We explain the absence of excluded volume by using a polymer melt, which contains 125 polymer chains each composed of 641 monomers (S9 Fig). The system was simulated considering only chain connectivity and Weeks-Chandler-Anderson type exclude volume interactions [69]. The latter manifests itself as the zero-valued probability of the bond length (*r*_*i,i*+1_) and intra-chain pairwise monomer distance (*r*_*ij*_) when *r <* 1 in units of the monomer diameter (the circular black dots in S9 Fig C and D). However, when the melt is coarse-grained such that we model 8 or 64 consecutive segments as a coarse-grained center, the coarse-grained segments overlap with each other. Together with the results shown in Fig 1B, it is justified to ignore the excluded volume and bending penalty beyond certain scales.

As a further test of the general applicability of HLM for the prediction of *n*-body contacts, we modeled the *β*-globin locus of mouse ES cells (S10 Fig) and the Pcdh*α* locus of mouse neural progenitor cells (S11 Fig) by using the Hi-C data reported by Bonev *at al* [70]. The pairwise and predicted triplet contact probabilites at multiple viewpoints are well correlated with the experiments (see detailed PC coefficient in the figure caption and S1 Table).

## Conclusions

To recapitulate, we have derived analytic expressions of *n*-body contact probabilities (for any *n* > 2) based on a recent chromatin polymer model (HLM) [50] and developed a method to predict multi-way chromatin contacts from Hi-C data.

First, the predicted triplet chromatin contacts are in reasonable correlation with the results of two independent measurements (i) Tri-C [17] and (ii) MC-4C [18], also in accordance with the genome-wide study using (iii) SPRITE [26]. Besides confirming the experimental findings, the suggested method was used to explore the multi-way chromatin contacts at any viewpoint of genomic locus of interest, which allowed us to discover some of key features not previously underscored in each measurement. (i) For the mouse *α*-globin locus, a cell-line dependent interaction pattern for the promoter is found when the viewpoint is anchored at the gene promoter. (ii) Although the previous study highlighted only the enrichment of long-range triplet contacts among CTCF binding sites in the ΔWAPL cells, we find that depletion of short-range contacts occurs for some triplets to compensate the enrichment. Our calculations also lend support to the aggregates of CTCF-bound boundary elements by explicitly showing the enrichments of four- and five-way chromatin contacts. (iii) Lastly, our analysis captures the enrichment of triplet contacts involving super-enhancers at transcriptionally active loci.

With an increasing contact order *n*, the probability of forming multi-way contacts decreases by orders of magnitude (see how the range of scale bars changes with increasing *n* in Fig 6A-C). Our theoretical approach can be used to circumvent this statistical limitation inherent to experimental detection of multi-way chromatin contacts. All the computer codes discussed here are provided in https://github.com/leiliu2015/HLM-Nbody, so that multi-way contacts can be calculated from an input Hi-C dataset. The methodology developed here will be of great help to elucidate the regulatory roles played by complex chromatin topology.

## Methods

### Numerical details for determining the stiffness matrix 𝒦*

Our previous numerical procedure [50, 51] to determine 𝒦*-matrix has been improved in this paper by adapting those in the recent modeling studies of chromatin by two other groups [71–73]. The best solution, 𝒦*, which is consistent with a given Hi-C data, was determined with Hi-C data by optimizing the cost function, 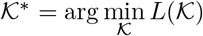 (see Fig 2). Different forms of the cost function can be conceived:

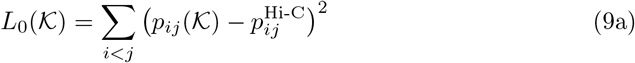

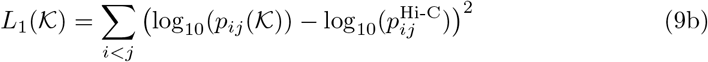

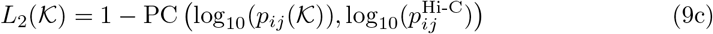

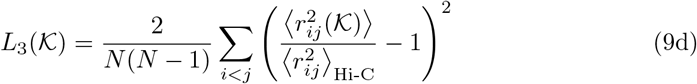

where 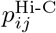 is the pairwise contact probability observed in Hi-C experiments, PC(*x, y*) stands for the Pearson correlation coefficient between *x* and *y*, and 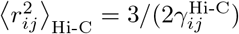 can be derived from 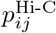 based on either Eq. 12 for Gaussian contact probability or Eq. S1 for contact probability in the form of Heaviside step function.

To avoid non-physical negative eigenvalues of the Kirchhoff matrix, the interaction strengths were previously required to be non-negative (*k*_*ij*_ ≥ 0) during the minimization [50, 71]. Shinkai *et al*. have overcome the issue of negative *k*_*ij*_ by alternately updating the backbone and non-backbone interaction strengths [72]. Even though there are some pairs with *k*_*ij*_ *<* 0, as long as the resulting 𝒦-matrix is positive-definite [72], the potential of mean force between the *i*- and *j*-th monomer is proportional to − ln *P*(*r*_*ij*_), which has a physically meaningful minimum at 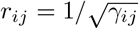.

We compared four different methods to optimize the cost functions: the first method minimizes the cost by random sampling (RS) of the parameter space [72]; the second one updates 𝒦 via steepest gradient descent (GD) [71]; the third and fourth methods, RMSprop [74] and ADAM [75], calculate both the gradient and its second moment to accelerate the model training [76].

As a case study, the decay of the cost function during the model training of a 2.4-Mb region on chr8 in mouse embryonic stem cells is shown in S12A Fig. The ADAM optimizer outperforms other methods significantly. Quality of the final model was calibrated by using the stratum adjusted correlation (SCC) [62] and PC between log_10_(*p*_*ij*_) from Hi-C and that from the model. As shown in the accompanying S2 Table, while *L*_1_ and *L*_3_ are both good candidates, *L*_2_ seems to be the best choice. However, the dependence of contact probability on the subchain size *s*, defined as 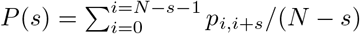, shows that the optimal model based on minimizing *L*_2_ underestimates the overall contacts (S12B Fig), although the pattern of log_10_(*p*_*ij*_) from the model is highly correlated with that from Hi-C. Similar conclusions can be drawn from training a model of a 10-Mb region on chr5 in GM12878 cells (S13 Fig), which our previous work used as a test case [50]. The models trained with new schemes clearly achieve better quality (S13 Fig D).

Taken together, in this study we calculated the 𝒦-matrix based on Hi-C data by minimizing *L*_1_ with *F*_0_, or minimizing *L*_3_ with *F*_1_ using ADAM optimizer, where *F*_0_ and *F*_1_ are two possible functional forms of cross-linking probability explained below.

### Derivations of the multi-body contact probability

Despite many potential biases in Hi-C [13], the frequency of two chromatin fragments being cross-linked in millions of cells is ideally determined by the probability density of their spatial distance *P*(*r*), and the efficiency of the cross-linking agent. The latter contribution can be included by the *r*-dependent cross-linking probability of fragments *F*(*r*). One can consider using a Gaussian function, 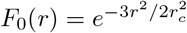, for cross-linking probability, which makes many-body contacts easier to handle mathematically. Alternatively, the Heaviside step function *F*_1_(*r*) = Θ(*r*_*c*_ − *r*) is conceived as well, as a natural choice of the cross-linking probability, such that fragment pairs are cross-linked if the spatial distance *r* is smaller than an effective capture radius of the cross-linking agent (*r*_*c*_).

The contact probabilities based on the two cross-linking probabilities, *F*_0_(*r*) and *F*_1_(*r*) are denoted by “*p*^(0)^” and “*p*^(1)^”, respectively. Unless stated otherwise, the results shown in the main text were calculated with *F*_0_(*r*), and “*p*” was used as a shorthand notation of “*p*^(0)^” for brevity.

Here we present the derivation of *n*-body contact probability based on the Gaussian cross-linking probability, *F*_0_(*r*). The derivation based on the Heaviside step function, *F*_1_(*r*), is provided in S1 Appendix.

### Generic many-body contact probability

Based on Eqs. 1 and 2, if we set, 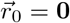, the many-body contact probability among *n* (≥ 2) monomers in a given set 𝒜 = (*i, j*, …) can be directly calculated as

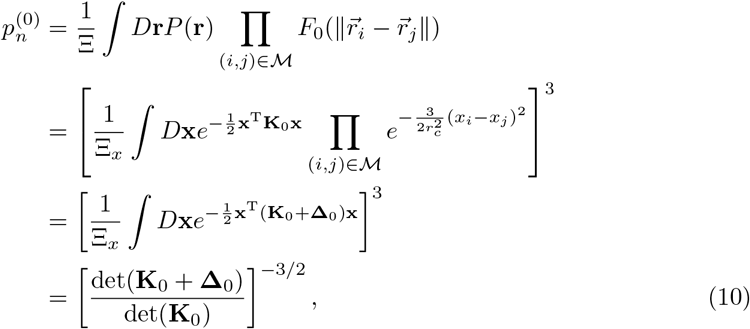

where *D***r** ≡ Π_*i*_ *d***r**_*i*_, *P* (**r**) is the probability density of the chain configuration (**r**) and M is the enumeration of all the pair-wise combinations of the elements in 𝒜. **Δ** is a matrix of elements **Δ**_*uv*_ whose values are given by

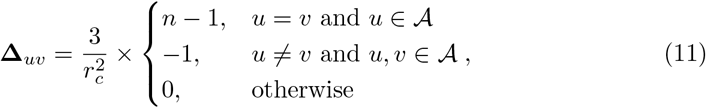

in which *u, v* ∈ (0, 1, 2, …, *N* − 1). The **K** and **Δ** with subscript “0” in Eq. 10 signifies the matrices whose 0-th row and column are removed from the original ones.

### Pairwise contact probability

To avoid computing determinants of large matrices in the general solution (Eqs. 10 and 11), *p*_*n*_ can alternatively be calculated based on *n*-body correlation function. For example, considering the probability density of the pairwise distance between the *i*-th and *j*-th monomer in three-dimensional space (Eq. 3), the pairwise contact probability between the *i*-th and *j*-th monomers can be formulated as [71, 72]

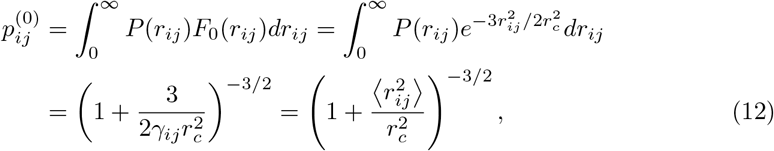

where Eq. 3 and 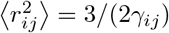 is used in the second row.

### Three-body contact probability

For three-body contact probability *p*_*ijk*_, we first consider the probability density of the distances between the *i*-th and *j*-th monomers, and between the *j*-th and *k*-th monomers projected on one dimension

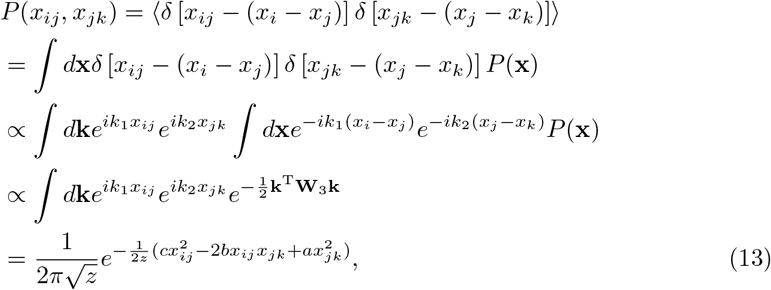

where **k** = (*k*_1_, *k*_2_), 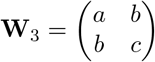 is a matrix with elements

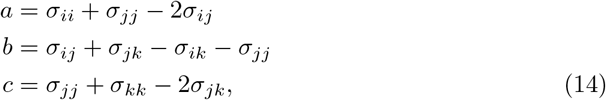

and

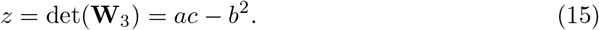

Remember that *σ* is the element of **K**^−1^, and *σ*_*ij*_ = 0 if *i* × *j* = 0. Then, the three-body contact probability in 3D is given by

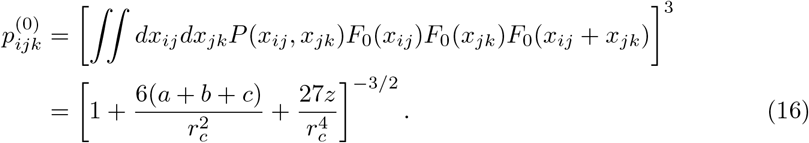

### Four-body contact probability

Four-body contact probability *p*_*ijkl*_ can be derived in the same manner. By defining **u**_*x*_ = (*x*_*ij*_, *x*_*jk*_, *x*_*kl*_)^T^, the one dimensional probability density of three pairwise distances, *P*(*x*_*ij*_, *x*_*jk*_, *x*_*kl*_), is given by

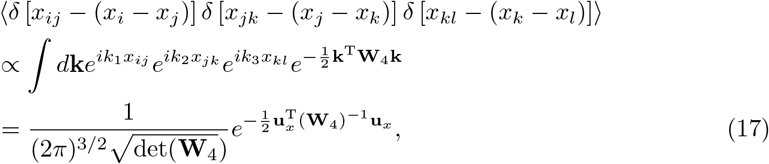

where now **k** = (*k*_1_, *k*_2_, *k*_3_). **W**_4_ has a matrix form of

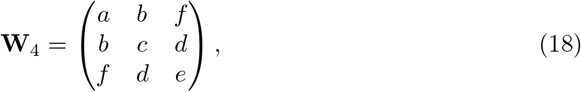

with elements in Eq. 14 and

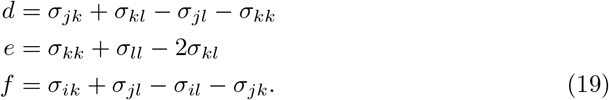

Then, from Eq. 17, one gets

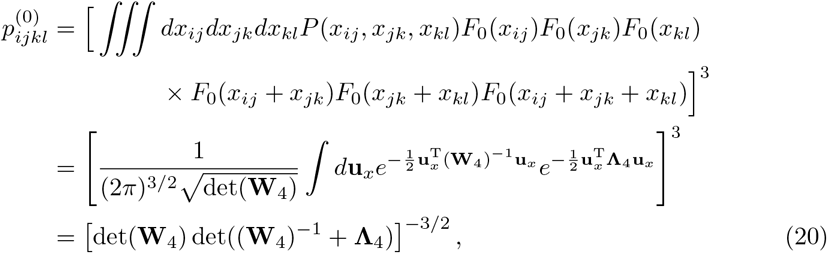

where

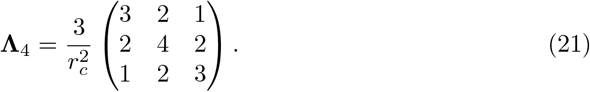

### Conditional pairwise contact probability

With super-resolution tracing, Bintu *et al*. examined whether the association between two chromatin loci facilitates or prevents the contact with a third locus [14]. More specifically, given three monomers of indices *i < j < k*, two conditional probabilities *p*(*r*_*jk*_ ≤ *r*_*c*_ | *r*_*ij*_ ≤ *r*_*c*_) and *p*(*r*_*jk*_ ≤ *r*_*c*_ | *r*_*ij*_ > *r*_*c*_) were compared with the marginal one *p*(*r*_*jk*_ ≤ *r*_*c*_) (see Fig 5 in Ref. [14]). They found “cooperativity”, namely *p*(*r*_*jk*_ ≤*r*_*c*_ | *r*_*ij*_ > *r*_*c*_) *< p*(*r*_*jk*_ ≤ *r*_*c*_) *< p*(*r*_*jk*_ ≤ *r*_*c*_ | *r*_*ij*_ ≤ *r*_*c*_), among more than 80% triplets of CTCF-bound and generic loci, as well as little effect of cohesin depletion induced by auxin treatment.

Assuming *F*_0_(*r*), we next aim at deriving the conditional contact probabilities of the *j*- and *k*-th monomers given that the *j*-th monomer is in contact with the *i*-th monomer 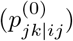, or given that the *j*-th monomer is in contact with the *i*-th monomer 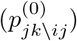, with an ordering of indices *i < j < k*. The first conditional probability can be calculated based on the relation that 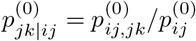, in which

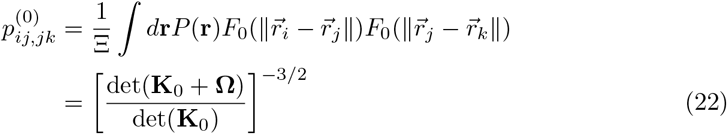

with non-zero elements 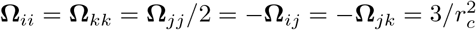, or equivalently

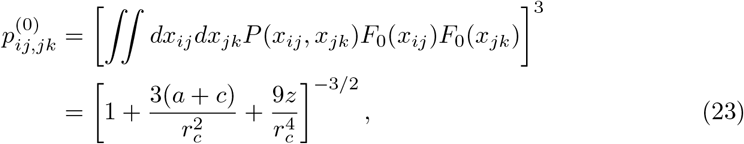

where we have applied Eq. 13 and 14. By using the relation that 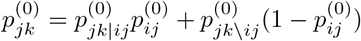, the second conditional pairwise contact probability can be determined as

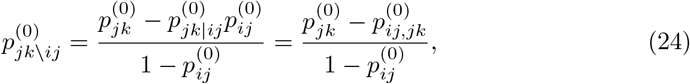

where the marginal contact probabilities 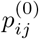 and 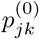 are given by Eq. 12.

Together with 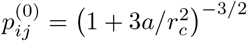 and 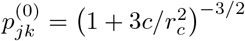, it is straightforward to prove that for any triplet

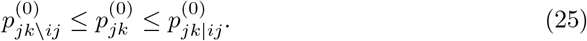

The equality signs hold if and only if *b* = 0 (see Eq.14), which corresponds to the case that *x*_*ij*_ and *x*_*jk*_ are completely uncorrelated.

### Gaussian polymer chain

At last we consider the simplest model, a Gaussian polymer chain, in which *k*_*ij*_ = *k*_0_ if |*i* − *j*| = 1 and *k*_*ij*_ = 0 otherwise. It follows that *σ*_*ij*_ = min(*i, j*)*/k*_0_. With Eq. 16, it straightforwardly leads to

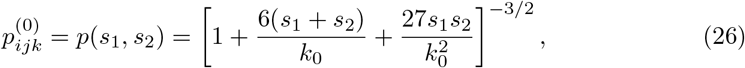

where *s*_1_ = *j* − *i* and *s*_2_ = *k* − *j* for any triplet with *i < j < k*.

There is a power-law decay of 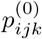 with *s*_1_ as 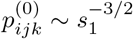 if *s*_1_ ≫ *s*_2_. It is also easy to prove that the quantity, *p*_*ijk*_*/p*_*ij*_*p*_*jk*_, discussed in Fig 6D is a nontrivial function of *s*_1_ and *s*_2_. If *s*_1_ + *s*_2_ = constant, it has its minimum at *s*_1_ = *s*_2_, and maximum at *s*_1_ = 1 or *s*_2_ = 1.

In addition, as a result of *b* = 0 in Eq. 14, we notice that 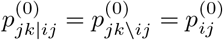 for a Gaussian chain (see Eq. 22-25).

## Code availability

The software package and associated documentation are available online (https://github.com/leiliu2015/HLM-Nbody),

## Supporting information

**S1 Appendix. Derivation of n-body contact probability based on the cross-linking probability modeled with the Heaviside step function**.

**S1 Fig. Stratified Pearson correlations of the HLM-predicted conact probabilities**. Stratified PC at *α*-globin locus of mouse **(A)** ES and **(B)** erythroid cells compared with Capture-C (2-body) and Tri-C (3-body) experiments.

**S2 Fig. Changes of Tri-C triplet contact frequencies in the** *α***-globin region of mouse erythroid cells with respect to ES cells**. The analysis was done at the viewpoints of R2 (top) and HS-39 (bottom). Following the statistical analysis in the experiment [17], we use the symbol * to mark all triplet interactions with significant changes (*P <* 0.01). The erythroid cell-specific regulatory hub and diffuse interactions among CTCF boundary sites, which are highlighted by dashed triangles, are both captured by Tri-C and our theory (Fig 3E).

**S3 Fig. Analysis of the changes in triplet contacts** 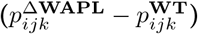 **and in pairwise contacts** 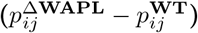 **among CTCF binding sites in WAPL lacking cells. (A)** The triplet contacts from MC-4C data [18] with respect to the viewpoints of E (top) and K (bottom). The triplet contacts predicted by HLM are shown in Fig 4D. **(B)** The enrichment of pairwise contacts between long-range CTCF binding sites (off-diagonal elements in red corresponding to 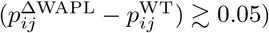 is counteracted by the depletion of contacts in the domains flanked by short-range CTCF binding sites (matrix elements along the diagonal block in blue corresponding to 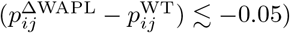. The data of pairwise contacts are obtained from Hi-C data [63].

**S4 Fig. WAPL depletion-induced fold changes of** *n***-body contact probability**. Fold changes of **(A)** four-body contacts double-anchored at sites E, K and **(B)** five-body contacts triple-anchored at sites E, H and K. The absolute change of contact probabilities is shown in Fig 4F and G, respectively.

**S5 Fig. Comparison between HLM and SPRITE**. Comparison between HLM and SPRITE similarly to those of Fig 5 in a 0.52 Mb region on human chr12. **(C)** From the viewpoint of the promoter of active ETV6 gene which encodes an ETS family transcription factor.

**S6 Fig. Many-body contacts of a Gaussian polymer chain calculated with step function**. Same as Fig 6 but using *F*_1_(*r*) (Heaviside step function) as the cross-linking probability.

**S7 Fig. Conditional pairwise contact probability with either a Gaussian (left column) or Heaviside step (right column) cross-linking probability. (A)** Heatmaps of log_10_(*p*_*ij*_) in a 1.23-Mb region on chr21 of IMR90 cells from Hi-C and log_10_(*p*_*ij*_) from HLM. **(B)** Comparison between the unconditioned contact probability *p*_*jk*_, the conditional contact probability, *p*_*jk*|*ij*_, and *p*_*jk\ij*_, calculated for CTCF-site triplets. **(C)** *p*_*jk*_, *p*_*jk*|*ij*_, and *p*_*jk\ij*_ calculated for all triplets *ijk* of *i < j < k*, which are sorted in an ascending order of *p*_*jk*_.

**S8 Fig. Triplet contacts predicted by simple rules with reference to Fig 3C and D. (A)** Three-body contact matrix at *α*-globin locus of mouse erythroid cells measured by Tri-C experiment or predicted by using three simple rules **(B-D). (E)** Pearson correlations between the results from 4 methods and the experiment.

**S9 Fig. Excluded volume in a polymer melt at different levels of coarse graining. (A)** A typical configuration of a dense polymer melt and **(B)** one polymer chain in the melt at three levels of coarse graining. The beads in **(B)** are colored differently along the chain, with a diameter of the most probable bond length at the corresponding scales. **(C)** Probability of the bond length and **(D)** intra-chain pairwise monomer distance.

**S10 Fig. HLM of mouse** *β***-globin locus on chr7 at resolution of 8kb. (A)** Pairwise contact probability from Hi-C [70] compared with HLM, which has a PC of 0.996. **(B)** Triplet contact probabilities from Tri-C compared with HLM, which are anchored at 3’HS1 and HS2 with PCs of 0.62 and 0.88, respectively.

**S11 Fig. HLM of mouse Pcdh***α* **locus on chr18 at 5 kb resolution (see also the caption of Fig S10)**. Compared with the MC-4C dataset, **(A)** the pairwise contact probability has a PC of 0.98, and **(B)** the triplet contact probabilities have PCs of 0.84, 0.76, 0.76, 0.88, and 0.70 at the viewpoint of Pcdh*α*1, Pcdh*α*11, Pcdh*α*c1, HS7, and HS5-1, respectively.

**S12 Fig. Comparison between the model trainings with different choices of cost functions and optimizers**. We trained a polymer model of a 2.4-Mb genomic region on chr8 in mouse ES cells at 25-kb resolution [70]. **(A)** The trajectories of various cost functions *L* in a log-log scale, minimized by using one of the four methods (RS, GD, RMSprop, and ADAM) with different cross-linking probabilities *F*_*α*_ (*α* = 0, 1). **(B)** Comparing *P*(*s*) from Hi-C and from three models, which were all trained with ADAM using *F*_0_, but with different forms of the cost functions. **(C)** Comparison of log_10_(*p*_*ij*_) from Hi-C (top) with that from the model trained by minimizing *L*_1_ with ADAM (bottom).

**S13 Fig. Comparison between the models trained for a 10-Mb genomic region at 50-kb resolution in GM12878 cells [77] with different choices of cost functions and optimizers. (A-C)** Same as the caption of S12 Fig. **(D)** Pearson correlations between Hi-C and HLM in our previous work [50] (the black line), and new models trained in this work (the colored lines) as a function of genomic separation, *s*.

**S1 Table. Genomic regions simulated in this work**.

**S2 Table. Pearson correlation (PC), stratum-adjusted correlation (SCC8) and distance-corrected Pearson correlation (DCPC9) of the contact probabilities predicted by SBS and HLM compared with Capture-C1 and Tri-C2 experiments**.

**S3 Table. Stratum adjusted correlation (SCC) and PC coefficients between Hi-C and HLM model**.

## Acknowledgments

We thank Dr. Yonghyun Song for careful reading of the manuscript and helpful comments. We thank the Center for Advanced Computation in KIAS for providing computing resources.

