## Supplementary Text, Figures for "Extracting multi-way chromatin contacts from Hi-C data"

### S1 Appendix

Derivation of $n$-body contact probability based on the cross-linking probability modeled with the Heaviside step function.

#### Pairwise contact probability

Along with Eq. 9, the pairwise contact probability assuming $F_1(r)$ can be determined by [1]

$$p_{ij}^{(1)} = \int_0^\infty P(r_{ij})\Theta(r_c - r_{ij})dr_{ij} = \int_0^{r_c} P(r_{ij})dr_{ij} = \text{erf}(\gamma_{ij}^{1/2} r_c) - 2r_c\sqrt{\frac{\gamma_{ij}}{\pi}} e^{-\gamma_{ij} r_c^2}, \tag{S1}$$

with $\text{erf}(x) = \frac{2}{\sqrt{\pi}} \int_0^x dte^{-t^2}$.

#### Three-body contact probability

The simultaneous contact probability among three sites $i$, $j$, and $k$ assuming $F_1$, $p_{ijk}^{(1)}$, can be defined as

$$p_{ijk}^{(1)} \equiv p\left((r_{ij} \leq r_c) \wedge (r_{ik} \leq r_c) \wedge (r_{jk} \leq r_c)\right) = \int_0^{r_c} p\left((r_{ik} \leq r_c) \wedge (r_{jk} \leq r_c) \mid r_{ij}\right) P(r_{ij})dr_{ij}, \tag{S2}$$

where $p\left((r_{ik} \leq r_c) \wedge (r_{jk} \leq r_c) \mid r_{ij}\right)$ $(\equiv p_{ik,jk|r_{ij}}^{(1)})$ stands for the probability of the $k$-th monomer being simultaneously in contact with other two monomers conditioned with the distance $r_{ij}$, which we evaluated using cylindrical coordinates $(\rho, \phi, z)$. Under the conditions of $\vec{r}_i = (0, 0, r_{ij}/2)$ and $\vec{r}_j = (0, 0, -r_{ij}/2)$, the position of $k$-th monomer is still described in terms of the Gaussian distribution

$$P(\vec{r}_k) = Ce^{-\gamma_{ij,k}(z_k - z_{ij,k}^*)^2} e^{-\gamma_{ij,k}\rho_k^2}, \tag{S3}$$

with the variance of the distribution $\gamma_{ij,k} = \det(\mathbf{K}_{ij})/\det(\mathbf{K}_{ijk})$, and a normalization constant $C = (\pi/\gamma_{ij,k})^{-3/2}$. $P(\vec{r}_k)$ is shifted along $z$-axis by

$$z_{ij,k}^* = \frac{r_{ij}}{2}\{(\mathbf{K}_{ij})^{-1} \cdot (\mathbf{k}_{\setminus\{i,j\}}^j - \mathbf{k}_{\setminus\{i,j\}}^i)\}_k, \tag{S4}$$

where $\mathbf{k}^m_{\backslash\{i,j\}}$ is the $m$-th column of the matrix $\mathbf{K}$ after removing the $i$-th and the $j$-th rows. For $r_{ij} \leq r_c$, the conditional probability $q_{ik,jk|r_{ij}}$ equals to an integral over the intersection formed between two spheres of a radius $r_c$ ,

$$
\begin{aligned}
p^{(1)}_{ik,jk|r_{ij}} &= \int_0^{2\pi} d\phi \int_{-D}^{D} dz \int_0^{R(z)} d\rho P(\vec{r}_k)\rho \\
&= C \int_0^{2\pi} d\phi \int_{-D}^{D} dz e^{-\gamma_{ij,k}(z-z^*_{ij,k})^2} \int_0^{R(z)} d\rho e^{-\gamma_{ij,k}\rho^2} \rho \\
&= -\sqrt{\frac{\gamma_{ij,k}}{\pi}} \int_{-D}^{D} dz e^{-\gamma_{ij,k}(z-z^*_{ij,k})^2} e^{-\gamma_{ij,k}\rho^2} \Big|_{\rho=0}^{\rho=R(z)} \\
&= \sqrt{\frac{\gamma_{ij,k}}{\pi}} \Big[ \int_{-D}^{D} dz e^{-\gamma_{ij,k}(z-z^*_{ij,k})^2} \\
&\quad - e^{-\gamma_{ij,k} r_c^2} \int_{-D}^{0} dz e^{-\gamma_{ij,k}(z-z^*_{ij,k})^2} e^{\gamma_{ij,k}(\frac{r_{ij}}{2}-z)^2} \\
&\quad - e^{-\gamma_{ij,k} r_c^2} \int_{0}^{D} dz e^{-\gamma_{ij,k}(z-z^*_{ij,k})^2} e^{\gamma_{ij,k}(\frac{r_{ij}}{2}+z)^2} \Big] \\
&= \sqrt{\frac{\gamma_{ij,k}}{\pi}} \Big[ I_0 + e^{-\gamma_{ij,k} r_c^2} I_- + e^{-\gamma_{ij,k} r_c^2} I_+ \Big],
\end{aligned}
\tag{S5}
$$

where $R(z) = (r_c^2 - (\frac{r_{ij}}{2} + |z|)^2)^{1/2}$ has been inserted in the third row. Considering that $D = r_c - r_{ij}/2$ ($> 0$), it is straightforward to calculate the integrals in the second to the last row of Eq. S5. One gets

$$
I_0 = \begin{cases} \frac{1}{2}\sqrt{\frac{\pi}{\gamma_{ij,k}}} \big[\mathrm{erf}(|z_-|) + \mathrm{erf}(|z_+|)\big], & z_- z_+ < 0 \\ \frac{1}{2}\sqrt{\frac{\pi}{\gamma_{ij,k}}} \big|\mathrm{erf}(|z_-|) - \mathrm{erf}(|z_+|)\big|, & z_- z_+ \geq 0 \end{cases},
\tag{S6}
$$

with $z_{\mp} = (\mp D - z^*_{ij,k})\gamma^{1/2}_{ij,k}$, and

$$
I_{\mp} = \begin{cases} -D, & r_{\mp} = 0 \\ \frac{e^{(\gamma_{ij,k} r_+ r_-)/4}}{\gamma_{ij,k} r_{\mp}} (1 - e^{\gamma_{ij,k} D r_{\mp}}), & r_{\mp} \neq 0 \end{cases},
\tag{S7}
$$

where $r_{\mp} = r_{ij} \pm 2z^*_{ij,k}$. The triplet contact probability in Eq. S2 can be computed numerically as a one-dimensional finite integral of a function of variable $r_{ij}$, together with Eqs. 9, S5-S7.

### Conditional pairwise contact probability

Similarly to the discussions in the **Subsection: Comparison with SPRITE** in the main text, the first conditional pairwise contact probability, $p(r_{jk} \leq r_c \mid r_{ij} \leq r_c)$, can be calculated based on the relation that $p(r_{jk} \leq r_c \mid r_{ij} \leq r_c) = p^{(1)}_{jk|r_{ij} \leq r_c} = p^{(1)}_{ij,jk}/p^{(1)}_{ij}$, where

$$
\begin{aligned}
p^{(1)}_{ij,jk} &= p((r_{jk} \leq r_c) \wedge (r_{ij} \leq r_c)) \\
&= \int_0^{r_c} P(r_{ij}) p(r_{jk} \leq r_c \mid r_{ij}) dr_{ij}.
\end{aligned}
\tag{S8}
$$

By setting $\vec{r}_i = (0, 0, r_{ij})$ and $\vec{r}_j = (0, 0, 0)$, we calculated $p(r_{jk} \leq r_c \mid r_{ij})$ ($\equiv p^{(1)}_{jk|r_{ij}}$), via a similar procedure of obtaining $p^{(1)}_{ik,jk|r_{ij}}$ (Eq.S5). The distribution of the $k$-th monomer, $P(\vec{r}_k)$, has the same form as Eq. S3, with the center being determined at $z^*_{ij,k} = r_{ij}\{(\mathbf{K}_{ij})^{-1} \cdot (-\mathbf{k}^i_{\backslash\{i,j\}})\}_k$. $p^{(1)}_{jk|r_{ij}}$ is evaluated by integrating over a sphere centered at the origin with a radius $r_c$,

$$\begin{aligned} p^{(1)}_{jk|r_{ij}} &= \int_0^{2\pi} d\phi \int_{-r_c}^{r_c} dz \int_0^{R(z)} d\rho P(\vec{r}_k)\rho \\ &= \sqrt{\frac{\gamma_{ij,k}}{\pi}}\Big[\int_{-r_c}^{r_c} dz e^{-\gamma_{ij,k}(z-z^*_{ij,k})^2} \\ &- \int_{-r_c}^{r_c} dz e^{-\gamma_{ij,k}(z-z^*_{ij,k})^2} e^{-\gamma_{ij,k}(r_c^2-z^2)}\Big] \\ &= \sqrt{\frac{\gamma_{ij,k}}{\pi}}\Big[I_0 - I_1\Big], \end{aligned} \tag{S9}$$

where $R(z) = (r_c^2 - z^2)^{1/2}$, $I_0$ is given in Eq. S6 with $z_{\mp} = (\mp r_c - z^*_{ij,k})\gamma^{1/2}_{ij,k}$, and

$$I_1 = \begin{cases} 2r_c e^{-\gamma_{ij,k}(z^{*2}_{ij,k}+r_c^2)}, & z^*_{ij,k} = 0 \\ \frac{e^{-\gamma_{ij,k}(z^{*2}_{ij,k}+r_c^2)}}{\gamma_{ij,k} z^*_{ij,k}} \sinh(2\gamma_{ij,k} z^*_{ij,k} r_c), & z^*_{ij,k} \neq 0 \end{cases}. \tag{S10}$$

After computing $p^{(1)}_{ij,jk}$ with Eq. S8-S9, the second conditional pairwise contact probability can be determined by

$$p^{(1)}_{jk|r_{ij}>r_c} = \frac{p^{(1)}_{jk} - p^{(1)}_{ij,jk}}{1 - p^{(1)}_{ij}}, \tag{S11}$$

where the marginal contact probabilities $p^{(1)}_{ij}$ and $p^{(1)}_{jk}$ are given by Eq. S1 in this case.

Note that neither $p^{(1)}_{jk|r_{ij}\leq r_c}$ nor $p^{(1)}_{ij,jk}$ equals to $p^{(1)}_{ijk}$ if $r_c > 0$. Although both loci are concurrently in contact with the $j$-th site, the contact between the $i$-th and $k$-th loci is not guaranteed. To be precise, what Bintu *et al.* have quantified [2] are not higher-order chromatin contacts.

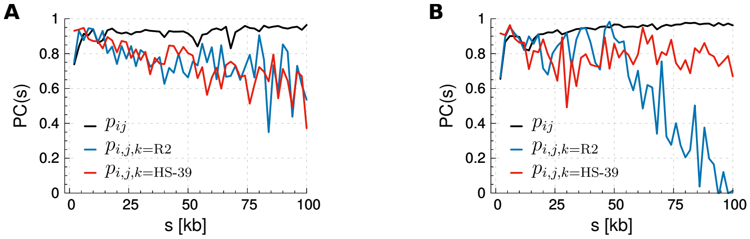


**Fig S1.** Stratified Pearson correlations of the HLM-predicted conact probabilities. Stratified PC at $\alpha$-globin locus of mouse **(A)** ES and **(B)** erythroid cells compared with Capture-C (2-body) and Tri-C (3-body) experiments.

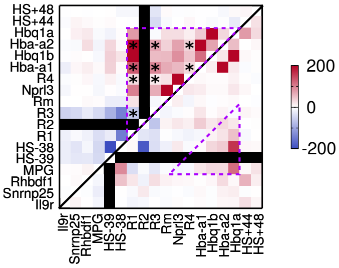


**Fig S2.** Changes of Tri-C triplet contact frequencies in the $\alpha$-globin region of mouse erythroid cells with respect to ES cells, calculated at the viewpoints of R2 (top) and HS-39 (bottom). Following the statistical analysis in the experiment [3], we use the symbol $*$ to mark all triplet interactions with significant changes ($P < 0.01$). The erythroid cell-specific regulatory hub and diffuse interactions among CTCF boundary sites, which are highlighted by dashed triangles, are both captured by Tri-C and our theory (Fig. 3E).

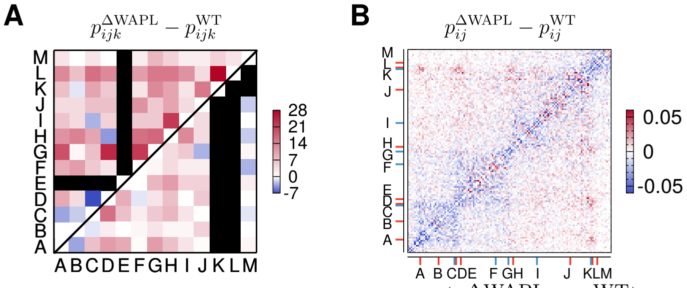


**Fig S3.** Analysis of the changes in triplet contacts ($p_{ijk}^{\Delta\mathrm{WAPL}} - p_{ijk}^{\mathrm{WT}}$) and in pairwise contacts ($p_{ij}^{\Delta\mathrm{WAPL}} - p_{ij}^{\mathrm{WT}}$) among CTCF binding sites in WAPL lacking cells. **(A)** The triplet contacts from MC-4C data [4] with respect to the viewpoints of E (top) and K (bottom). The triplet contacts predicted by HLM are shown on the left panel of Fig. 4D. **(B)** The enrichment of pairwise contacts between long-range CTCF binding sites (off-diagonal elements in red corresponding to $(p_{ij}^{\Delta\mathrm{WAPL}} - p_{ij}^{\mathrm{WT}}) \gtrsim 0.05$) is counteracted by the depletion of contacts in the domains flanked by short-range CTCF binding sites (matrix elements along the diagonal block in blue corresponding to $(p_{ij}^{\Delta\mathrm{WAPL}} - p_{ij}^{\mathrm{WT}}) \lesssim -0.05$). The data of pairwise contacts are obtained from Hi-C data [5].

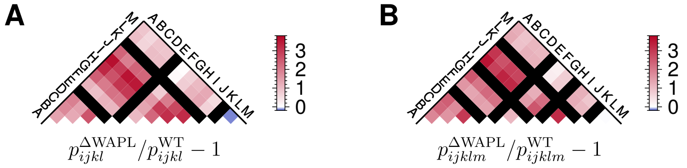


**Fig S4.** Fold changes of **(A)** four-body contacts double-anchored at sites E, K and **(B)** five-body contacts triple-anchored at sites E, H and K induced by WAPL depletion. The absolute change of contact probabilities is shown in Fig. 4F and G, respectively.

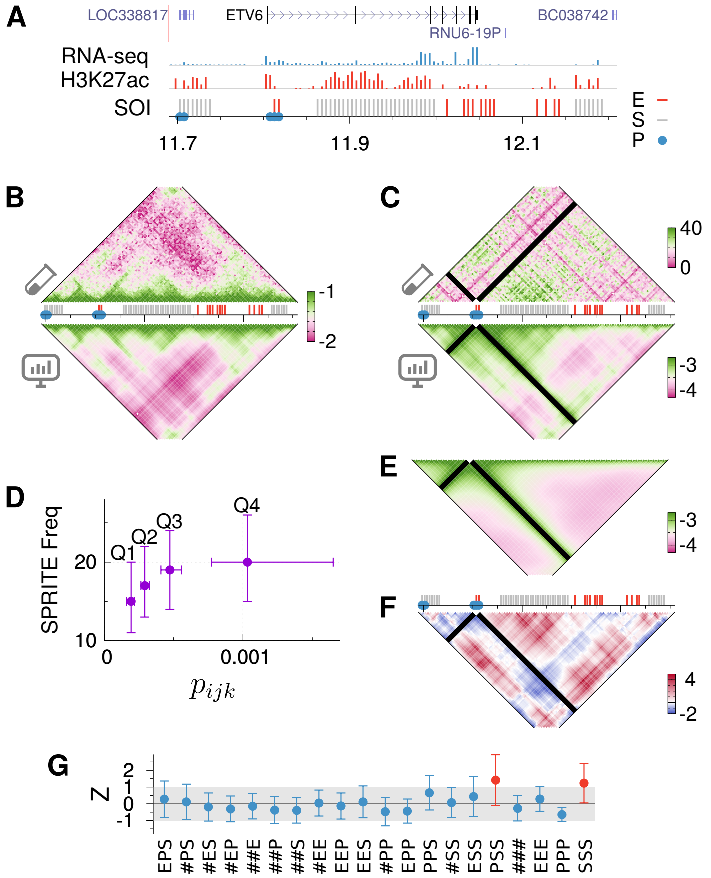


**Fig S5.** Similar comparison between HLM and SPRITE as Fig. 5 in a 0.52 Mb region on human chr12. **(C)** From the viewpoint of the promoter of active ETV6 gene which encodes an ETS family transcription factor.

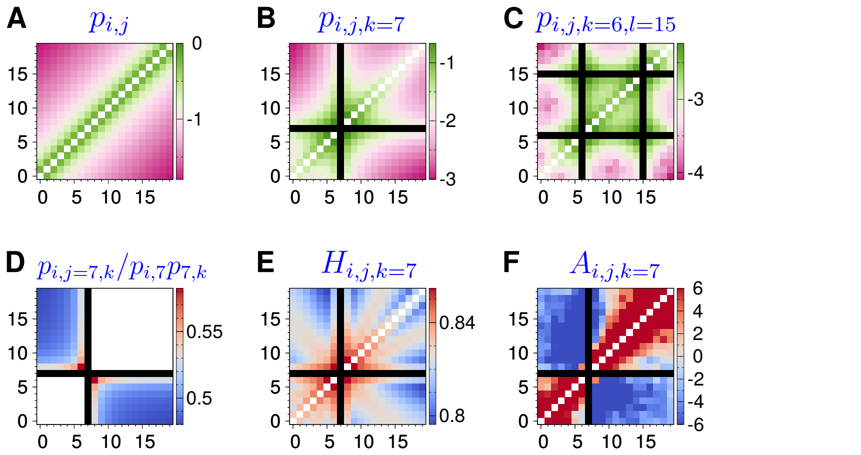


**Fig S6.** Same as Fig. 6 but using $F_1(r)$ (Heaviside step function) as the cross-linking probability.

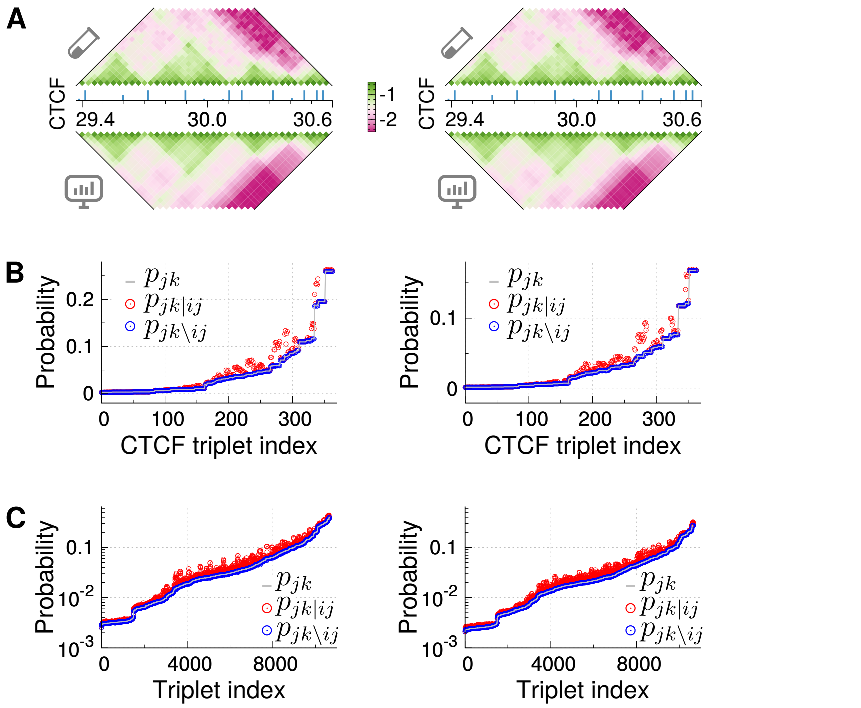


**Fig S7.** Conditional pairwise contact probability with either a Gaussian (left column) or Heaviside step (right column) cross-linking probability. **(A)** Heatmaps of $\log_{10}(p_{ij})$ in a 1.23-Mb region on chr21 of IMR90 cells from Hi-C and $\log_{10}(p_{ij})$ from HLM. **(B)** Comparison between the unconditioned contact probability $p_{jk}$, the conditional contact probability, $p_{jk|ij}$, and $p_{jk\setminus ij}$, calculated for CTCF-site triplets. **(C)** $p_{jk}$, $p_{jk|ij}$, and $p_{jk\setminus ij}$ calculated for all triplets $ijk$ of $i < j < k$, which are sorted in an ascending order of $p_{jk}$.

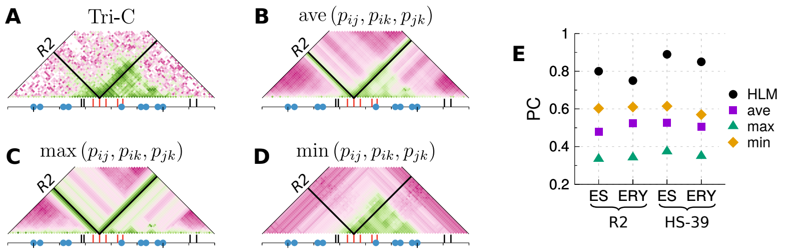


**Fig S8.** Triplet contacts predicted by simple rules with reference to Fig 3C and D. **(A)** Three-body contact matrix at $\alpha$-globin locus of mouse erythroid cells measured by Tri-C experiment or predicted by using three simple rules **(B-D)**. **(E)** Pearson correlations between the results from 4 methods and the experiment.

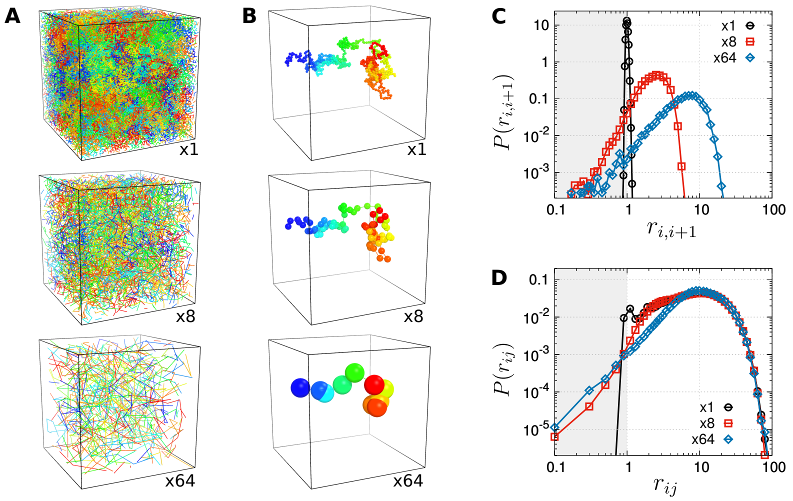


**Fig S9.** Excluded volume in a polymer melt at different levels of coarse graining. **(A)** A typical configuration of a dense polymer melt and **(B)** one polymer chain in the melt at three levels of coarse graining. The beads in **(B)** are colored differently along the chain, with a diameter of the most probable bond length at the corresponding scales. **(C)** Probability of the bond length and **(D)** intra-chain pairwise monomer distance.

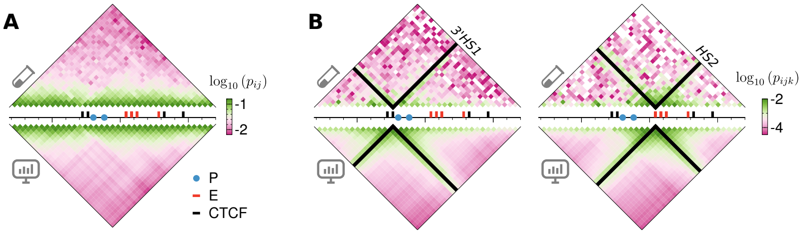


**Fig S10.** HLM of mouse $\beta$-globin locus on chr7 at resolution of 8kb. **(A)** Pairwise contact probability from Hi-C [6] compared with HLM, which has a PC of 0.996. **(B)** Triplet contact probabilities from Tri-C compared with HLM, which are anchored at 3'HS1 and HS2 with PCs of 0.62 and 0.88, respectively.

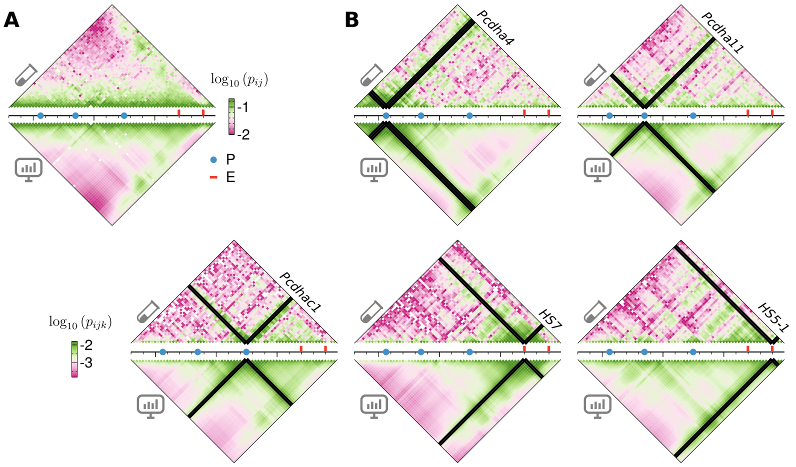


**Fig S11.** HLM of mouse Pcdh$\alpha$ locus on chr18 at 5 kb resolution (see also the caption of Fig S10). Compared with the MC-4C dataset, **(A)** the pairwise contact probability has a PC of 0.98, and **(B)** the triplet contact probabilities have PCs of 0.84, 0.76, 0.76, 0.88, and 0.70 at the viewpoint of Pcdh$\alpha$1, Pcdh$\alpha$11, Pcdh$\alpha$c1, HS1, and HS5-1, respectively.

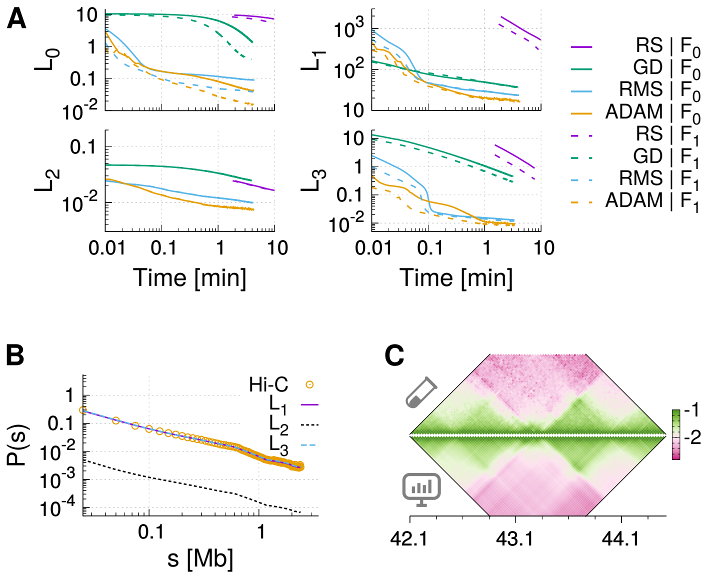


**Fig S12.** Comparison between different choices of the model training. We trained a polymer model of a 2.4-Mb genomic region on chr8 in mouse ES cells at 25-kb resolution [6]. **(A)** The trajectories of various cost functions $L$ in a log-log scale, minimized by using one of the four methods (RS, GD, RMSprop, and ADAM) with different cross-linking probabilities $F_\alpha$ ($\alpha = 0, 1$). **(B)** Comparing $P(s)$ from Hi-C and from three models, which were all trained with ADAM using $F_0$, but with different forms of the cost functions. **(C)** Comparison of $\log_{10}(p_{ij})$ from Hi-C (top) with that from the model trained by minimizing $L_1$ with ADAM (bottom).

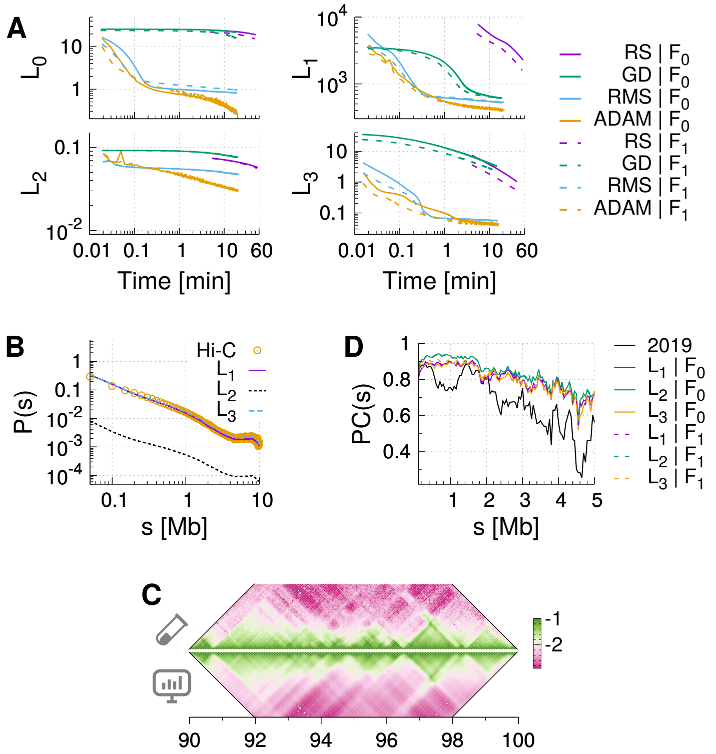


**Fig S13.** Comparison between different choices of the model trained for a 10-Mb genomic region at 50-kb resolution in GM12878 cells [7]. **(A-C)** Same as the caption of Fig. S12. **(D)** Pearson correlations between Hi-C and HLM in our previous work [1] (the black line), and new models trained in this work (the colored lines) as a function of genomic separation, $s$.
