## Supplementary Table for "Extracting multi-way chromatin contacts from Hi-C data"

TABLE S1. Genomic regions simulated in this work.

| Species | Chr | Region [a] | Res[b] | $N$ | Cell line | 2-body[c] | PC [d] | Viewpoint | 3-body[e] | PC [f] | Figure |
|---|---|---|---|---|---|---|---|---|---|---|---|
| mouse | 11 | 32,000,000-32,300,000 | 2 | 150 | ES | [1] | 0.99 | 32,151,060-32,151,883 | [2] | 0.80 | Fig. 3C |
| | | | | | | | | 32,137,176-32,137,426 | [2] | 0.89 | Fig. 3D |
| | | | | | ERY | [1] | 0.98 | 32,151,060-32,151,883 | [2] | 0.75 | Fig. 3C |
| | | | | | | | | 32,137,176-32,137,426 | [2] | 0.85 | Fig. 3D |
| | 8 | 120,800,000-122,080,000 | 10 | 128 | HAP1 WT | [3] | 0.97 | 121,126,666-121,145,791 | [4] | 0.83 | Fig. 4B |
| | | | | | | | | 121,941,808-121,960,933 | [4] | 0.71 | Fig. 4C |
| | | | | | HAP1 ΔWAPL | [3] | 0.96 | 121,126,666-121,145,791 | [4] | 0.81 | Fig. 4B |
| | | | | | | | | 121,941,808-121,960,933 | [4] | 0.73 | Fig. 4C |
| | 7 | 110,836,514-111,140,514 | 8 | 38 | ES | [5] | 1.00 | 110,955,508-110,955,666 | [2] | 0.62 | S10 Fig.B |
| | | | | | | | | 111,009,550-111,009,749 | [2] | 0.88 | S10 Fig.B |
| | 18 | 37,059,654-37,399,654 | 5 | 68 | NPC | [5] | 0.98 | 37,109,610-37,114,710 | [4] | 0.84 | S11 Fig.B |
| | | | | | | | | 37,167,503-37,172,603 | [4] | 0.76 | S11 Fig.B |
| | | | | | | | | 37,247,128-37,252,228 | [4] | 0.76 | S11 Fig.B |
| | | | | | | | | 37,335,329-37,340,429 | [4] | 0.88 | S11 Fig.B |
| | | | | | | | | 37,374,676-37,379,776 | [4] | 0.70 | S11 Fig.B |
| human | 18 | 60,675,000-61,120,000 | 5 | 89 | GM12878 | [6] | 0.99 | 60,980,000-60,990,000 | [7] | 0.18 | Fig. 5C |
| | 12 | 11,690,000-12,210,000 | 5 | 104 | GM12878 | [6] | 0.99 | 11,800,000-11,815,000 | [7] | 0.16 | S12 Fig.C |
| | 21 | 29,370,000-30,600,000 | 30 | 41 | IMR90 | [6] | 0.99 | | | | S13 Fig |

[a] The reference genome assemblies of mouse and human are mm9 and hg19, respectively.
[b] The model resolutions in units of kb.
[c] References of the experimental pairwise contacts datasets.
[d] Pearson correlation coefficient of two-body contact probabilities between Hi-C/Capture-C and HLM.
[e] References of the experimental triplet contacts datasets.
[f] Pearson correlation coefficient of three-body contact probabilities between 3-body experiments and HLM.

TABLE S2. Pearson correlation (PC), stratum-adjusted correlation (SCC[8]) and distance-corrected Pearson correlation (DCPC[9]) of the contact probabilities predicted by SBS[10] and HLM compared with Capture-C[1] and Tri-C[2] experiments ("NA" stands for not available).

| | | Pairwise contacts | | Triplet contacts at R2 | | Triplet contacts at HS-39 | |
|---|---|---|---|---|---|---|---|
| | | (SBS, Cap-C) | (HLM, Cap-C) | (SBS, Tri-C) | (HLM, Tri-C) | (SBS, Tri-C) | (HLM, Tri-C) |
| ES | PC | 0.96 | 0.99 | 0.80 | 0.80 | 0.84 | 0.89 |
| | SCC | 0.75 | 0.88 | 0.66 | 0.72 | 0.77 | 0.82 |
| | DCPC | 0.87 | 0.91 | NA | 0.68 | NA | 0.65 |
| ERY | PC | 0.96 | 0.98 | 0.80 | 0.75 | 0.77 | 0.85 |
| | SCC | 0.92 | 0.93 | 0.48 | 0.66 | 0.79 | 0.80 |
| | DCPC | 0.91 | 0.97 | NA | 0.70 | NA | 0.72 |

TABLE S3. Stratum adjusted correlation (SCC) and PC coefficients between Hi-C and HLM model, which was trained with different cross-linking probabilities, cost functions, and optimizers (see also Fig. S12). Each coefficient value is the averaged result of five independent trainings.

| | | | RS | GD | RMS | ADAM |
|---|---|---|---|---|---|---|
| SCC | $F_0$ | $L_0$ | -1.872e-02 | 5.684e-02 | 7.326e-01 | 7.250e-01 |
| | | $L_1$ | 7.700e-02 | 7.400e-01 | 7.936e-01 | **8.418e-01** |
| | | $L_2$ | 7.191e-01 | 5.811e-01 | 8.159e-01 | **8.537e-01** |
| | | $L_3$ | -1.872e-02 | 9.641e-02 | 7.960e-01 | 8.390e-01 |
| PC | $F_0$ | $L_0$ | 8.194e-01 | 9.378e-01 | 9.829e-01 | 9.835e-01 |
| | | $L_1$ | 8.731e-01 | 9.818e-01 | 9.884e-01 | **9.914e-01** |
| | | $L_2$ | 9.836e-01 | 9.753e-01 | 9.901e-01 | **9.924e-01** |
| | | $L_3$ | 8.1934-01 | 9.025e-01 | 9.883e-01 | 9.913e-01 |
| SCC | $F_1$ | $L_0$ | 6.318e-02 | 6.798e-02 | 7.661e-01 | 8.073e-01 |
| | | $L_1$ | 3.944e-01 | 7.488e-01 | 7.933e-01 | 8.441e-01 |
| | | $L_2$ | 7.161e-01 | 5.809e-01 | 8.155e-01 | **8.543e-01** |
| | | $L_3$ | 8.207e-02 | 1.040e-01 | 7.956e-01 | **8.454e-01** |
| PC | $F_1$ | $L_0$ | 8.561e-01 | 9.339e-01 | 9.843e-01 | 9.900e-01 |
| | | $L_1$ | 9.423e-01 | 9.825e-01 | 9.884e-01 | 9.917e-01 |
| | | $L_2$ | 9.834e-01 | 9.753e-01 | 9.901e-01 | **9.924e-01** |
| | | $L_3$ | 8.627e-01 | 9.003e-01 | 9.885e-01 | **9.918e-01** |
